# Microbial stir bars: light-activated rotation of tethered bacterial cells to enhance mixing in stagnant fluids

**DOI:** 10.1101/2023.01.26.525760

**Authors:** Jyoti P Gurung, Moein N Kashani, Charitha M de Silva, Matthew AB Baker

**Affiliations:** School of Biotechnology and Biomolecular Science, UNSW Sydney, Sydney, NSW, 2052, Australia; Future Industries Institute, University of South Australia, Mawson Lakes, SA, 5095, Australia; Australian National Fabrication Facility – South Australia Node, Mawson Lakes, SA, 5095, Australia; School of Mechanical and Manufacturing Engineering, UNSW Sydney, Sydney, NSW, 2052, Australia; ARC Centre of Excellence in Synthetic Biology, UNSW Sydney, Sydney, NSW, 2052, Australia

**Keywords:** μPIV, mixing, bacterial flagellar motor, fluorescence, Proteorhodopsin

## Abstract

Microfluidics devices are gaining significant interest in biomedical applications. However, in a micron-scale device, reaction speed is often limited by the slow rate of diffusion of the reagents. Several active and passive micro-mixers have been fabricated to enhance mixing in microfluidic devices. Here, we demonstrate external control of mixing by rotating a rodshaped bacterial cell. This rotation is driven by ion transit across the bacterial flagellar stator complex. We first measured the flow fields generated by rotating a single bacterial cell rotationally locked to rotate either clockwise (CW) or counterclockwise (CCW). Micro-Particle Image Velocimetry (μPIV) and Particle Tracking Velocimetry results showed that a bacterial cell of ~ 2.75 μm long, rotating at 5.75 ± 0.39 Hz in a counterclockwise direction could generate distinct micro-vortices with circular flow fields with a mean velocity of 4.72 ± 1.67 μm/s and maximum velocity of 7.90 μm/s in aqueous solution. We verified our experimental data with a numerical simulation at matched flow conditions which revealed vortices of similar dimensions and speed. We observed that the flow-field diminished with increasing z-height above the plane of the rotating cell. Lastly, we showed we could activate and tune rotational mixing remotely using strains engineered with Proteorhodopsin (PR), where rotation could be activated by controlled external illumination using green laser light (561 nm).

## Introduction

Magnetic stir bars are routinely used for mixing static solutions, that is, with no flow conditions, also termed stagnant fluids. These stir bars enhance mixing where solutions are vigorously advected within the container due to shearing induced by the circular motion of vortices (Halasz et al., 2007). In general, there are three mechanisms by which mixing is obtained: advection by shearing, convection by heating, and diffusion due to a concentration gradient. Microfluidics offers fine control of small reaction volumes at the micron scale. However, rapid mixing is still a challenge in microfluidic systems as these systems typically exist in conditions where viscous forces dominate over inertial forces and where Reynold’s number is low and often less than one. Because of the micron-sized channels in microfluidic devices, fluid flow is laminar, thus, mixing relies on the diffusion of molecules across the interface of fluids that are flowing parallel to each other. Such mixing is reliant on diffusion, which is typically slow in comparison with mixing by shearing and convection. Various strategies can be used to enhance mixing using convection and shearing to distribute large materials more vigorously than diffusion. In microfluidics, enhance mixing is obtained by applying external energy such as an electric field or a magnetic field (active mixing) or by shaping fluid flows using the structure of the device (passive mixing) (Lee et al., 2011). One example of passive mixing is the fabrication of serpentine-shaped channels, which creates a physical obstacle to the direction of flowing fluid, thus enhancing mixing by increasing the interfacial surface area of contact between laminar flow streams (Javaid et al., 2017).

Mixing in fluids that are not flowing, such as stagnant fluids trapped in micron-sized spaces requires energy to initiate fluid motion. Thus, active mixing by applied external fields is the primary method to induce mixing in these conditions. These fields, such as electric, acoustic, and magnetic fields can be controlled remotely and with specific energy and patterning. One example is the application of a magnetic field to rotate micron-sized magnetic microspheres to enhance mixing of stagnant fluids at the micron scale (Shanko et al., 2022). Although magnetic microspheres are efficient mixers, setting up magnets around a microfluidic device can be labour-intensive, micron-sized electromagnets are expensive to fabricate, and in some cases magnetic particles can be intrusive to biological cells or biomolecules (Shanko et al., 2019).

Here we sought to enhance mixing through the use of stir-bar shaped microorganisms such as bacteria. One advantage of using a bio-inspired mixing strategy is the capacity for biocompatibility and the ease of synthesis, at scale, leveraging genetic engineering tools and methods in synthetic biology (Carlsen & Sitti, 2014). Bacteria are mostly found in habitats which exhibit low Reynold’s number conditions, like the fluid conditions in microfluidic systems (Lauga, 2016). Despite the dominance of viscous force in low Reynolds number regimes, Bacteria use thin filamentous flagella (~10 μm in length) embedded in their cell membrane for locomotion. These flagella are driven by the rotation of a transmembrane nanometer-sized molecular machine known as a bacterial flagellar motor (BFM), which rotates with a maximum rotational speed up to 10,000 Hz (Sowa & Berry, 2008). The bacterial flagellar motor is an ideal candidate to drive mixing: it has high rotational speed and we have decades of understanding in how to genetically engineer the motor to change the rotational direction and sensitivity to different stimuli such as nutrients, light and magnetic fields (Gurung et al., 2020). The properties of bacterial motility and chemotactic sensing have been utilized to fabricate various microfluidics devices for mixing (M. J. Kim & Breuer, 2007), actuating the fluid flow (Al-Fandi et al., 2006), sensing pH (Krasnopeeva et al., 2020), and generating power (Tung & Kim, 2006).

In this paper, we used genetically engineered motile bacteria to mix stagnant fluids at the micron scale. Enhanced mixing driven by from surface-adhered cells rotating their flagellar bundle has been demonstrated in bacterial carpets (H. Kim et al., 2015). In our work, we focused on mixing driven by the rotation of a single motor driving the rotation of the rod-shaped bacterial cell body tethered to the surface by a single truncated filament (we term these ‘microbial stir bars’). To establish the potential of microbial stir bars as mixers, we measured strength and features of the flow-fields that were generated by rotation of a single cell in controlled conditions. Previous work used 1 μm polystyrene particles positioned near the edge of a rotating cell to track the flow induced by the rotation (Al-Fandi et al., 2010). However, more detailed characterization of flow field has not yet been performed. Here we quantified the flow field around single and multiple rotating cells and measured how this flow field evolved as a function of distance away from the plane of rotation.

Overall, we investigated the flow-field generated by the rotation of microbial stir bars using microscale particle image velocimetry (micro-PIV) in three spatial dimensions with higher spatial and temporal resolution than previously measured. We performed numerical simulations in matched geometries to compare with our experimental data and investigated flow fields generated by multiple cells rotating spatially close to each other. Lastly, we demonstrated simple external light-activation of mixing using genetically engineered bacteria which expressed the transmembrane light driven proton pump PR (Arlt et al., 2018).

## Results

### Directional control over cell rotation and particle tracing

Fluorescence microscopy was used for the micro-PIV experiments to investigate the flow field generated by a rotating bacterial cell in its surrounding aqueous solution (Figure 1a). Two types of green, fluorescent bacterial samples, one rotating in counterclockwise (CCW) and the other in clockwise (CW) were tethered suspended into an aqueous solution of fluorescent 200 nm beads (Figure 1c). Flow velocity was calculated by tracing the displacement of red fluorescent particles in a defined time interval induced by CCW and CW rotating cell (Figure 1c). The mean rotational speed of the CCW rotating cell was measured as 5.75 ± 0.39 Hz, and CW rotating cell was measured as 3.54 ± 0.58 Hz (Figure 1d). Both the CCW rotating cell and the CW rotating cell were approximately 2.5 long and covered the circular area of diameter ~ 3.32 μm and ~ 2.8 μm, respectively (Figure 1c, dotted circle; SI Figure 1). Rotational Reynolds number (Re) was quantified on the order of 10^-4^, given by the equation, Re = ωR^2^/ν for both cell types (where ‘ω’ is the angular velocity, ‘R’ is characteristic length, and ν is kinematic viscosity).

**Figure 1.**
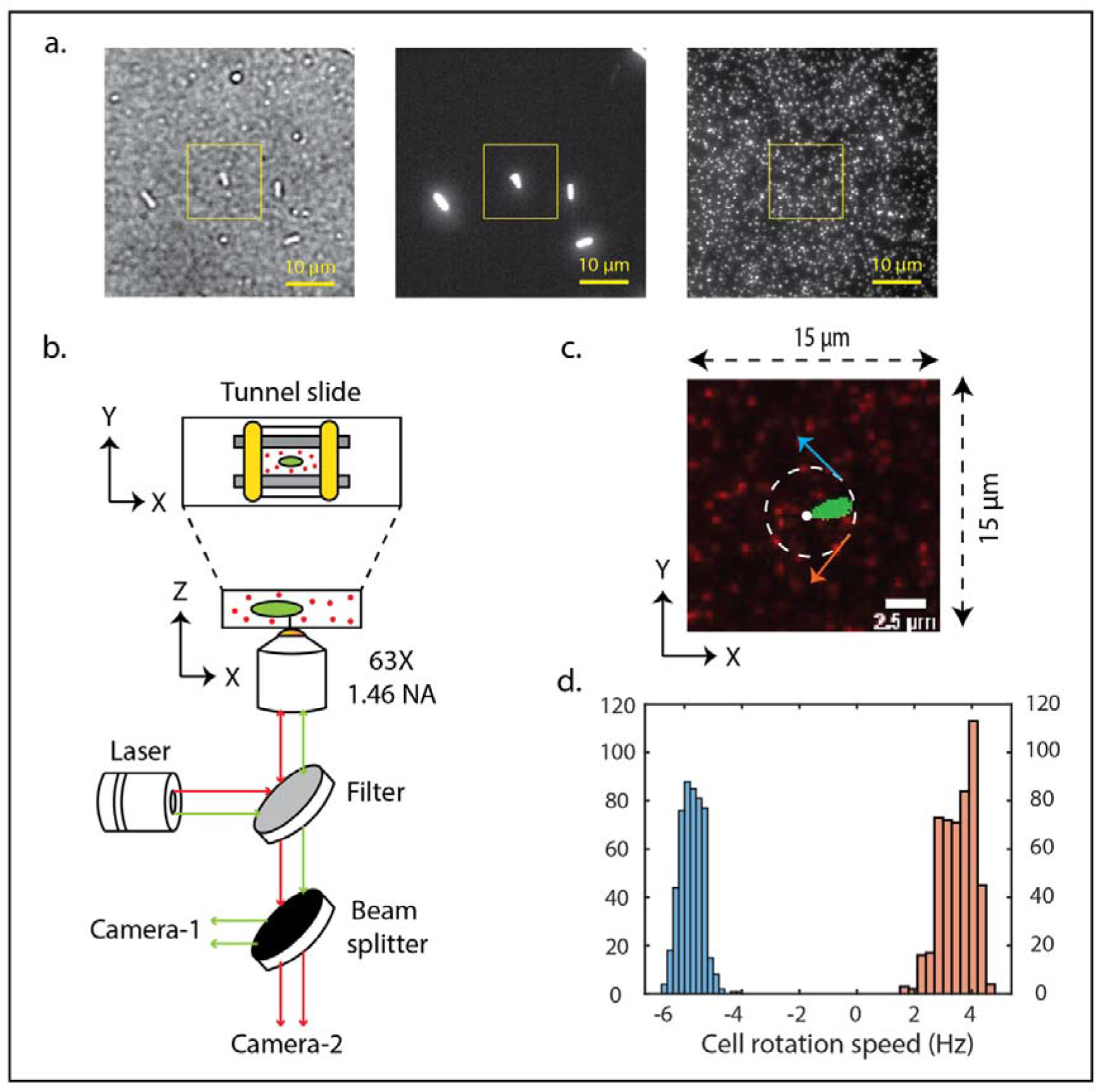
Experimental system used for micro-PIV to investigate flow fields generated by rotating bacterial cells. (a) Left – bright field image of bacterial cell and fluorescent tracer particles, middle – image of green, fluorescent bacterial cell acquired on Camera-1, and right – image of red, fluorescent tracer particles acquired at Camera-2. Yellow-outlined box represents the region of interest for analysis (15 by 15 μm). (b) Schematic of fluorescence microscope used for the micro-PIV experiment. Tunnel slide (top view) was loaded with bacterial cell (green) and fluorescent tracer particles (red) for micro-PIV, sealed with coverslip sealant (yellow). (c) Fluorescent image showing cells (green) and microspheres (red). Rotational direction CCW and CW indicated by blue and orange arrow respectively. (d) Histogram of cell rotational speed showing distribution of cell rotation speed in Hz for CCW (negative Hz, blue) and CW (positive Hz, orange) rotating cell.

### Flow fields generated by single cell rotation

For the CCW rotating single cell, a flow field with distinct pattern and directionality was observed near the region where the cell was rotating. The flow directionality, represented by velocity vector arrows (left image, Fig. 2a) and the velocity magnitude (middle image, Figure 2a), was visualized to examine the penetration of the flow field into the solution away from the boundary of the cell. Vorticity was measured to calculate the rotational circulation of fluid (right image Fig. 2a). A single cell rotating in CCW direction with the speed of 5.75 ± 0.39 Hz generated the distinct micro-vortices with mean velocity of 4.72 ± 1.67 μm/s, maximum velocity of 7.90 μm/s and maximum vorticity of 12.7 s^-1^.

**Figure 2.**
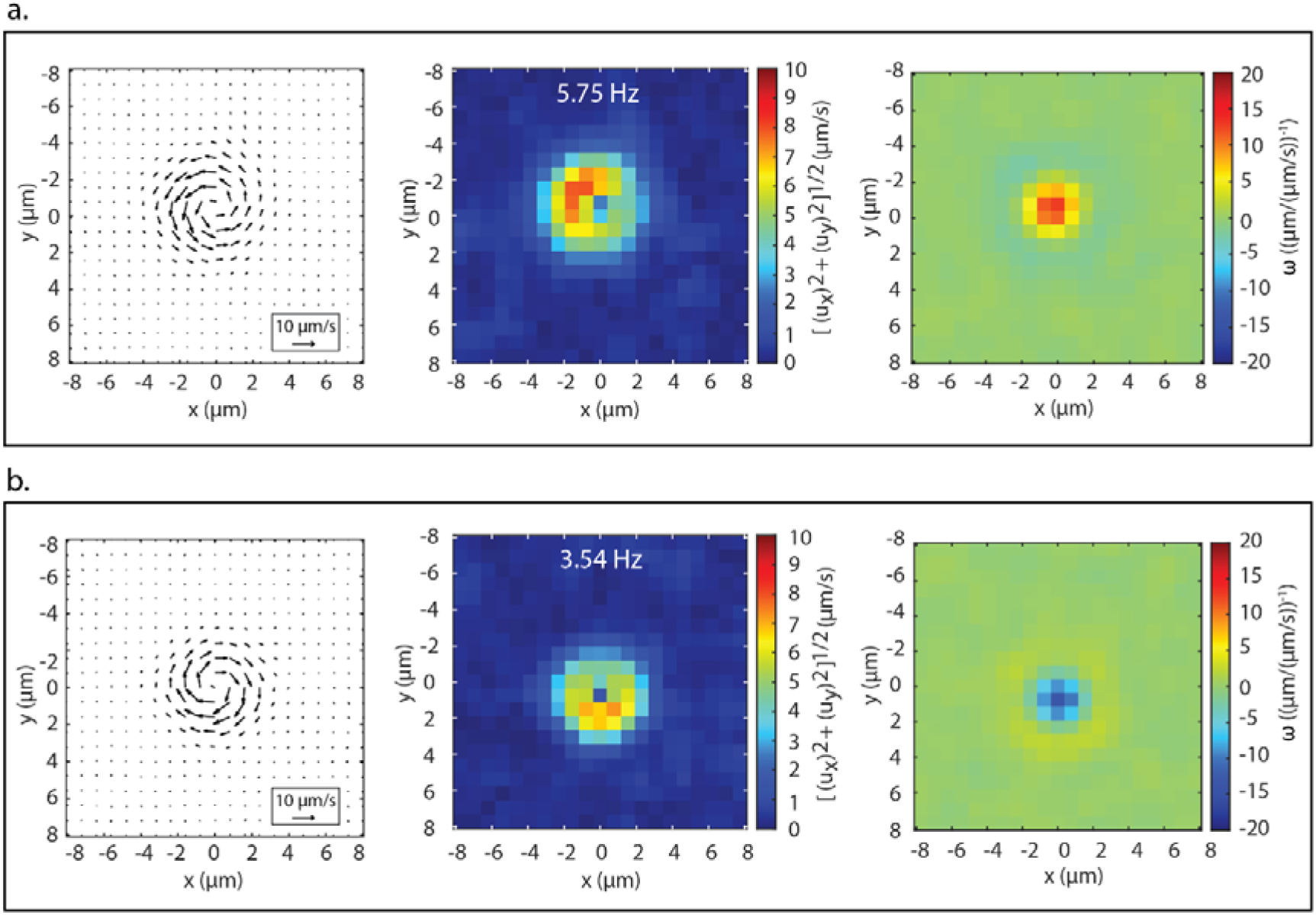
Micro-PIV results of flow fields generated by single cell rotation. (a) CCW rotating cell at the speed of 5.75±0.39 Hz across x-y plane (16 by 16 μm). (b) CW rotating cell at the speed of 3.54±0.58 Hz across x-y plane (16 by 16 μm, velocity calculated over individual PIV pixels of size 1.5 μm x 1.5 μm). Velocity arrow (left), velocity magnitude (middle), and vorticity (right).

Likewise for the opposite direction, the CW cell rotated with a speed of 3.54 ± 0.58 Hz generating a vortex with opposite directionality (Figure 2b). A single cell rotating in CW direction with the speed of 3.54 ± 0.58 Hz generated mean velocity of 3.95 ± 1.72 μm/s with maximum velocity of 7.26 μm/s and maximum vorticity of ~ 13.7 s^-1^. Velocity magnitude and vorticity increased with increasing speed for both CCW and CW rotating cells (Figure 3a & 3b). Simple linear regression fitting showed that velocity magnitude is linearly correlated to the speed of CCW and CW rotating cell with R^2^ value of 0.83 and 0.96 respectively whereas vorticity did not show significant linear relationship with the rotational speed (SI Figure 2a). Mean velocity at each time point was measured across the region of interest in the surrounding fluid to determine the time series evolution of velocity generated by rotating cell (SI Figure 3). For two selected regions of interest of the same size, the mean velocity across all PIV pixels for the CCW (5.75 ± 0.39 Hz) and CW (3.54 ± 0.58 Hz) rotating cell was greater in magnitude in the region with rotating cell than in the region without any rotating cells (SI Figure 3).

**Figure 3.**
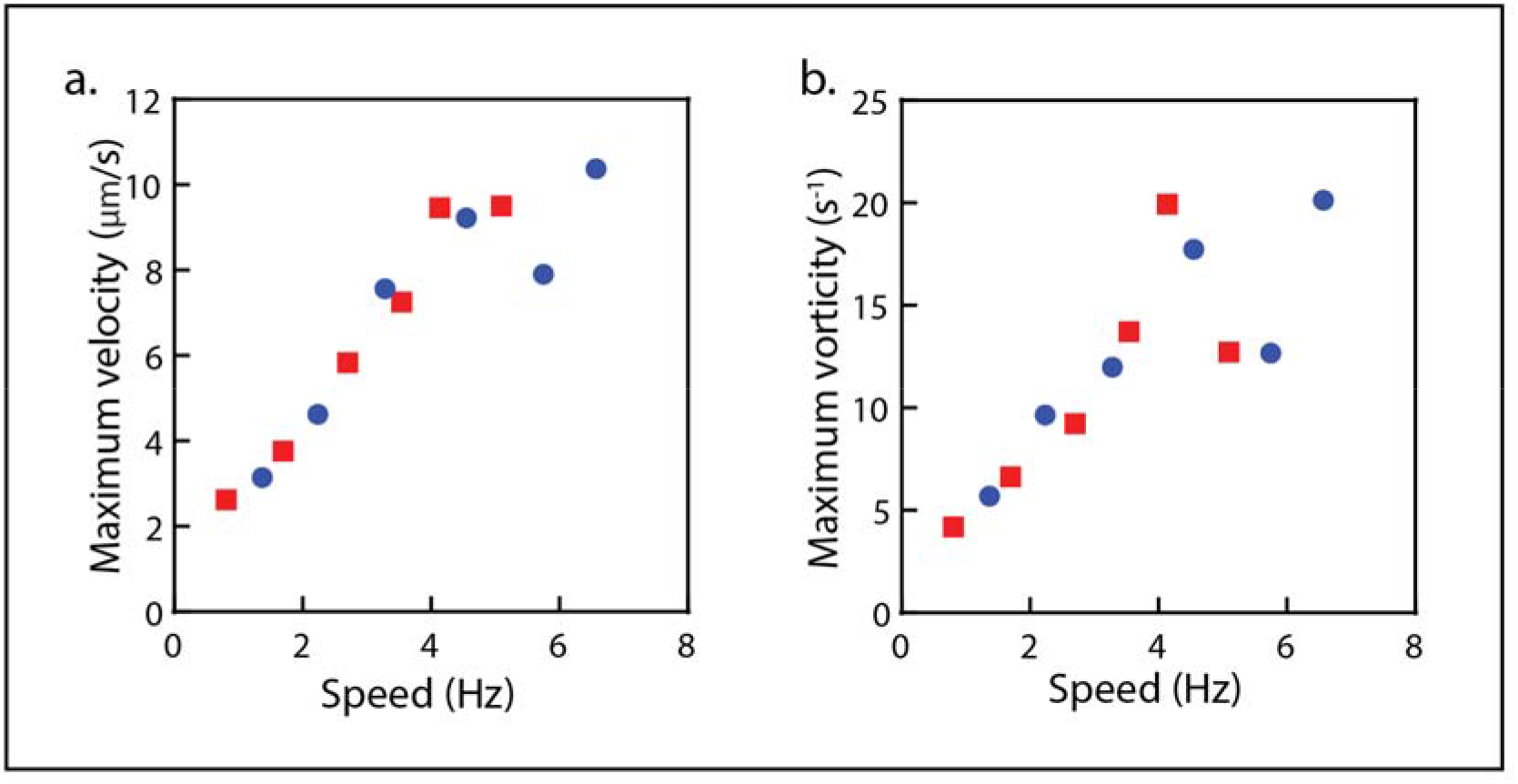
Blue circle/line represents ‘CCW rotating cell’, red square/line represents ‘CW rotating cell’, and black line represents ‘no rotating cell’. (a) Maximum fluid velocity taken over all PIV pixels (each occupying a region of 1.5μm x 1.5 μm i.e., equivalent to 16 x 16 pixel) for CCW and CW rotating cells (same cells as Fig. 2a and Fig. 2b respectively, with different speeds). (b) Maximum vorticity generated by CCW and CW rotating cell with different speeds.

### Flow fields generated at different z-heights

After confirming generation of vortices as a measure of circular flow pattern for both CCW and CW rotating cells, we measured azimuthal velocity (or tangential velocity) to determine the magnitude of flow velocity generated across the plane of cell rotation i.e., x-y plane. Azimuthal velocity was measured for angles between 0 and 2π in the x-y plane vs radius, where radius was defined as a distance from the center of rotation (SI Figure 4). For both CCW (5.75 ± 0.39 Hz) and CW (3.54 ± 0.58 Hz) rotating cells, velocity magnitude decreased with the increasing distance away from the center of rotation (Figure 4a). Velocity decreased exponentially with distance from the center of rotation for both CCW and CW cell types (R^2^ value of 0.99 and 0.98 respectively; SI Figure 2b).

**Figure 4.**
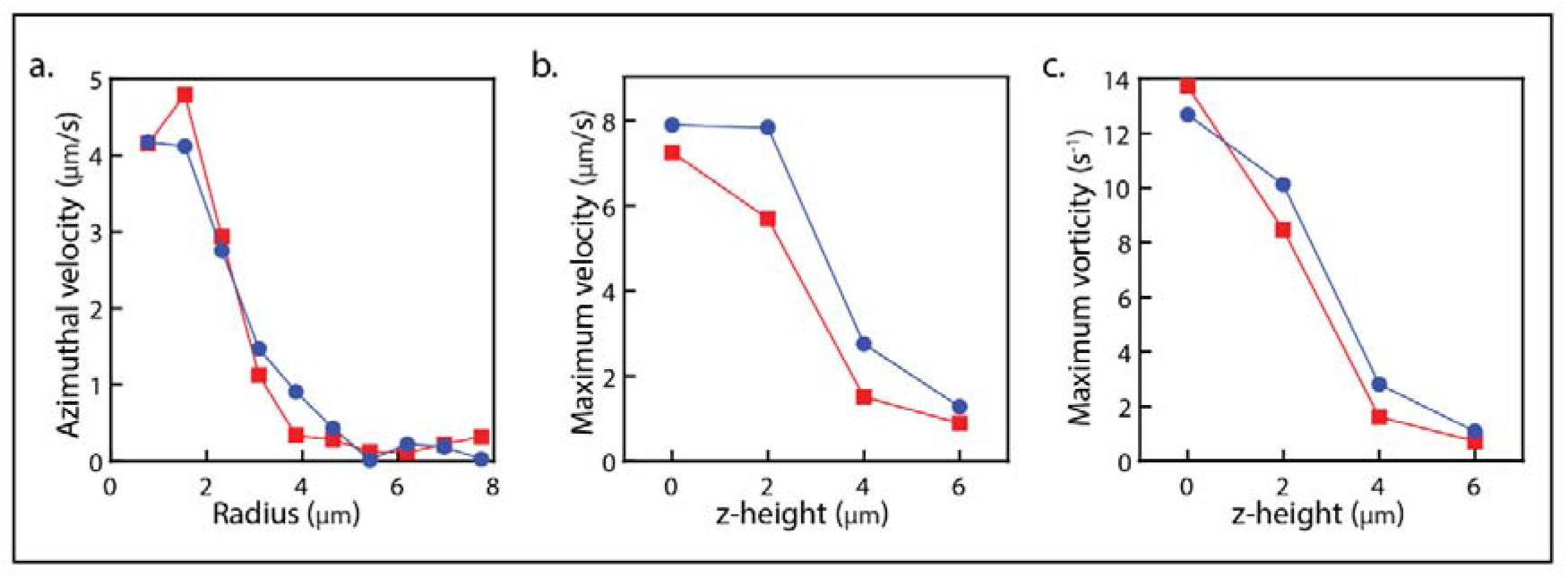
Blue circle/line represents ‘CCW rotating cell’ and red square/line represents ‘CW rotating cell’. (a) Azimuthal/tangential velocity generated by CCW and CW rotating cells at the distance away from the center of rotation (radius) across x-y plane. (b) Maximum velocity generated by CCW and CW rotating cells at different z-heights. (c) Maximum vorticity generated by CCW and CW rotating cells at different z-heights.

After analyzing the velocity profile in the x-y spatial plane, we investigated the flow field along the z-height to understand the penetration of the flow field away from the surface and into the chamber. Using the Hi-Lo (highly inclined and laminated optical sheet) mode of illumination (Tokunaga et al., 2008), we could illuminate and track our particles away from the surface at even increments of 2 μm. This increments of 2 μm was defined based on a calculation of depth of field for our microscope (Meinhart et al., 2000). The flow field generated by the CCW rotating cell with the speed of 5.75 ± 0.39 Hz was prominent until a z-height of 4 μm in comparison with the CW rotating cell with the speed of 3.54 ± 0.58 Hz where the flow field was not visible at 4 μm (Figure 4b and 5). In comparison, the flow field was not observed and detected at the z-height of 6 μm for both type of cells. Like the velocity, which exponentially decreased with increasing distance away from the center of the rotation in the x-y plane, we also observed an exponential decrease of velocity with increasing z-height along the z plane for both CCW and CW cell types (R^2^ value of 0.98 and 0.97 respectively (SI zure 2b).

**Figure 5.**
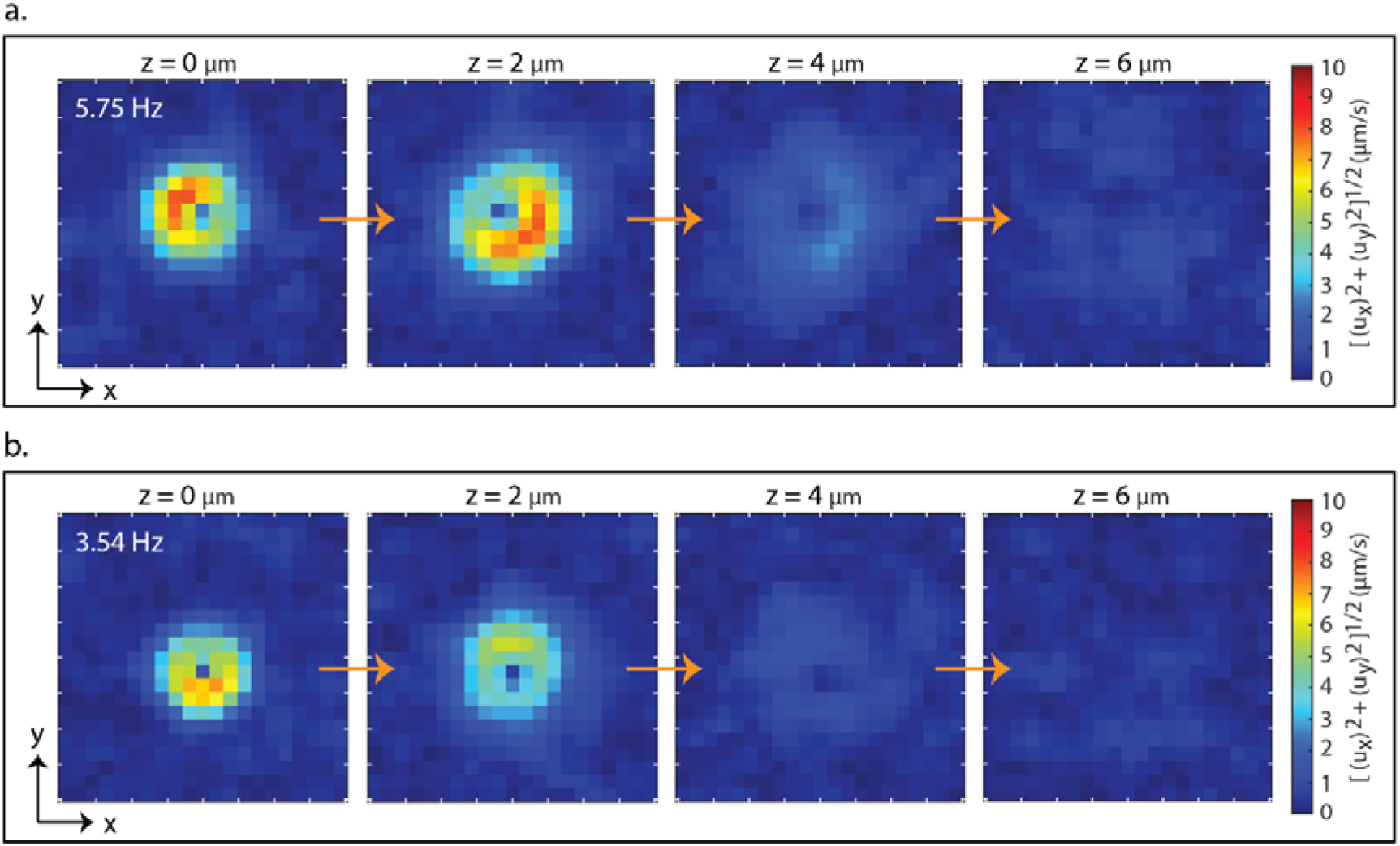
Time-averaged velocity generated by single-cell rotation at different z-heights. (a) CCW rotating cells at the speed of 5.75 ± 0.39 Hz across the x-y plane (16 by 16 μm) at 0, 2, 4, and 6 μm in z-height (left to right). (b) CW rotating cells at the speed of 3.54 ± 0.58 Hz across the x-y plane (16 by 16 μm) at 0, 2, 4, and 6 μm in z-height (left to right).

### Numerical simulation

Computation Fluid Dynamics (CFD) simulations were performed to investigate the flow field generated by rotating cells on the surrounding fluid till steady state conditions were reached (detailed description in Materials and Methods). Cells were defined as rods 2.5 μm long and 1.0 μm wide, capped with hemispheres, surrounded by the fluid with viscosity defined as 1 centipoise (cP) (close to water at 20°C). Parameters such as cell-speed (5.75 Hz for CCW rotating cell and 3.54 Hz for CW rotating cell) and x-y planes at different z-height (up to 6 μm) were defined to perform simulations of cells which matched our experimental conditions (SI Figures 5 & 6). Simulated data of velocity magnitude were generated at each time point with time step of 0.001 seconds till the cell completed one full rotation for both CCW and CW rotating cells (Supplementary Movies 2-9).

As per our experimental observations (Figure 5), the velocity flow field decreased with increasing z-heights for both CCW and CW rotating cells (Figure 6). For both CCW and CW rotating cells, exponential decay of velocity on increasing z-heights from 0 μm to 6 μm was observed with R^2^ value of 0.99 (SI Figure 5). In the x-y plane of the cell rotation, velocity decreased with increasing distance away from the edge of the cell for both CCW and CW rotating cells (Figure 6). For CCW cell rotating with the speed of 5.75 HZ, maximum velocity across the cell region was 100 μm/s, 11 μm/s, 4.4 μm/s, and 3.3 μm/s at the z-heights of 0 μm, 2 μm, 4 μm, and 6 μm respectively. Similarly, for CW cell rotating at the speed of 3.54 Hz, maximum velocity was 61 μm/s, 6.6 μm/s, 2.7 μm/s, and 2.0 μm/s at the z-heights of 0 μm, 2 μm, 4 μm, and 6 μm respectively. In this measure, simulated velocity of 100 μm/s for CCW rotating cell was in close agreement to the experiment where maximum velocity was 71.96 μm/s at the z-height of 0 μm. Likewise for the CW cell, maximum rotational velocity from simulation was 61 μm/s was in near agreement with maximum rotational velocity of 56.11 μm/s from the experiment at the z-height of 0 μm (SI Figure 7).

**Figure 6.**
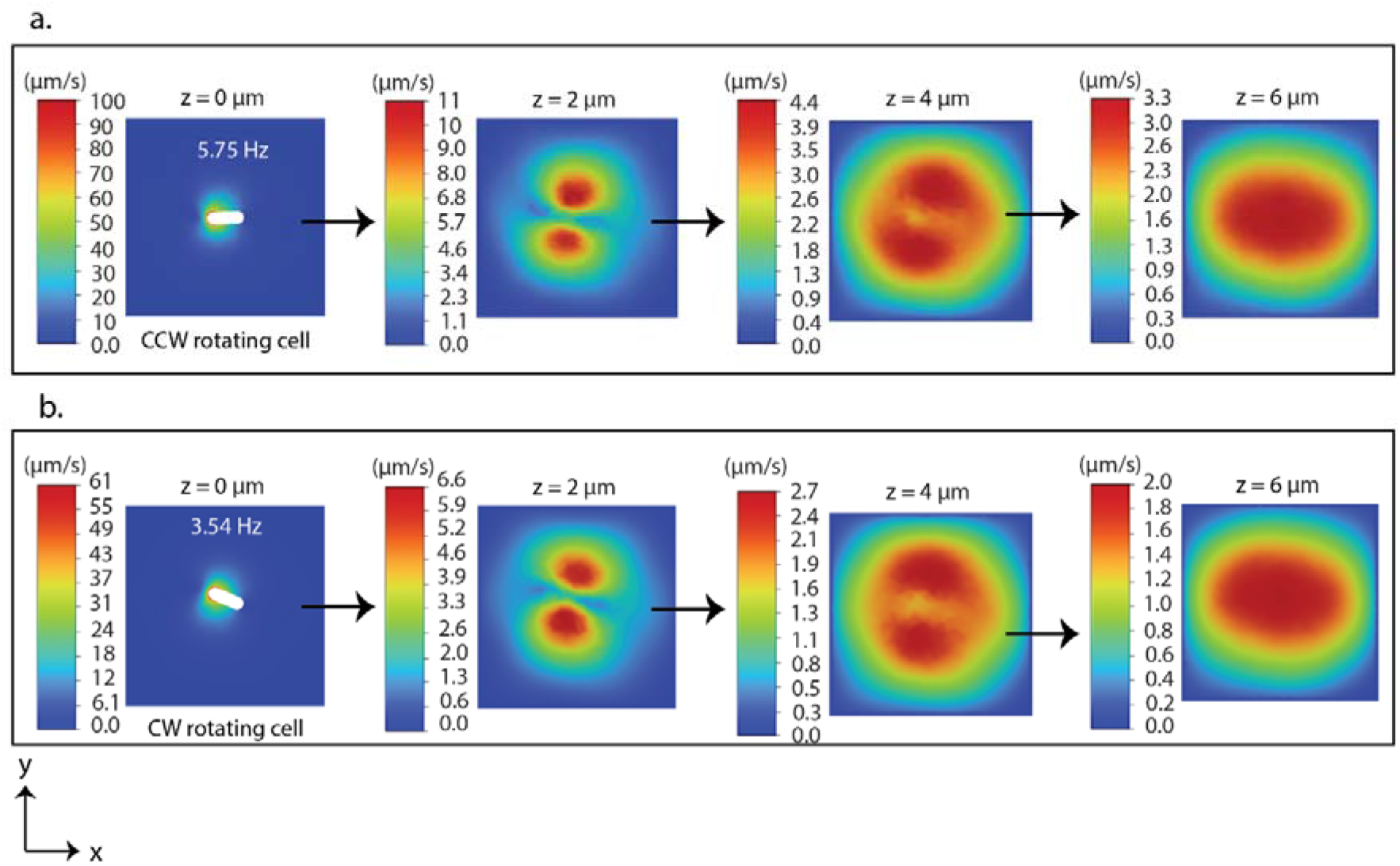
Numerical simulation of instantaneous velocity magnitude generated by single-cell rotation at 0, 2, 4, and 6 μm in z-height. (a) CCW rotating cell with the speed of 5.75 Hz across the x-y plane (15 by 15 μm) at 0.175 seconds. (b) CW rotating cell with the speed of 3.54 Hz across the x-y plane (15 by 15 μm) at 0.301 seconds. Both cells were elongated in rod like shape with axis of rotation at the edge, defined by the length of 2.75 μm and width of 1.0 μm. Direct comparison of experiment with simulation is shown in SI Figure 9.

To test for possible confounding surface effects, we also simulated the CCW rotating cell (1.13 Hz) in the middle of a region of interest 5 um from the bottom surface and top of the chamber. In these simulations the velocity decayed by 70% within 1 um from the rotating plane of the cell symmetrically in both directions (SI Figure 8).

### Flow fields generated by multiple cell rotation

To examine the potential interaction between adjacent cells’ flow fields, we examined the flow field of two cells in proximity. Two CCW cells rotating at the mean speed of 2.45 ± 2.23 Hz and 4.50 ± 1.48 Hz were separated by 10.47 μm (Figure 7a). Each of these cells generated a distinct velocity profile and microvortices. These micro vortices were isolated at this separation distance with no apparent hydrodynamic interaction (Figure 7a). Similarly, two CW cells rotating at the mean speed of 2.66 ± 1.16 Hz and 4.10 ± 0.64 Hz, which were also separated at the distance of 9.24 μm generated microvortices that showed no apparent hydrodynamic interaction (Figure 7b). The flow field diminished with increasing z-heights for both CCW and CW rotating multiple cells where the flow pattern was not apparent beyond a z-height of 4 μm (SI Figure 10). Furthermore, we examined the flow field for the multiple cells (more than two cells) rotating adjacent to each other. We observed microvortices for individual cells, but no distinguishable interaction in the region among multiple cells (SI Figure 11).

**Figure 7.**
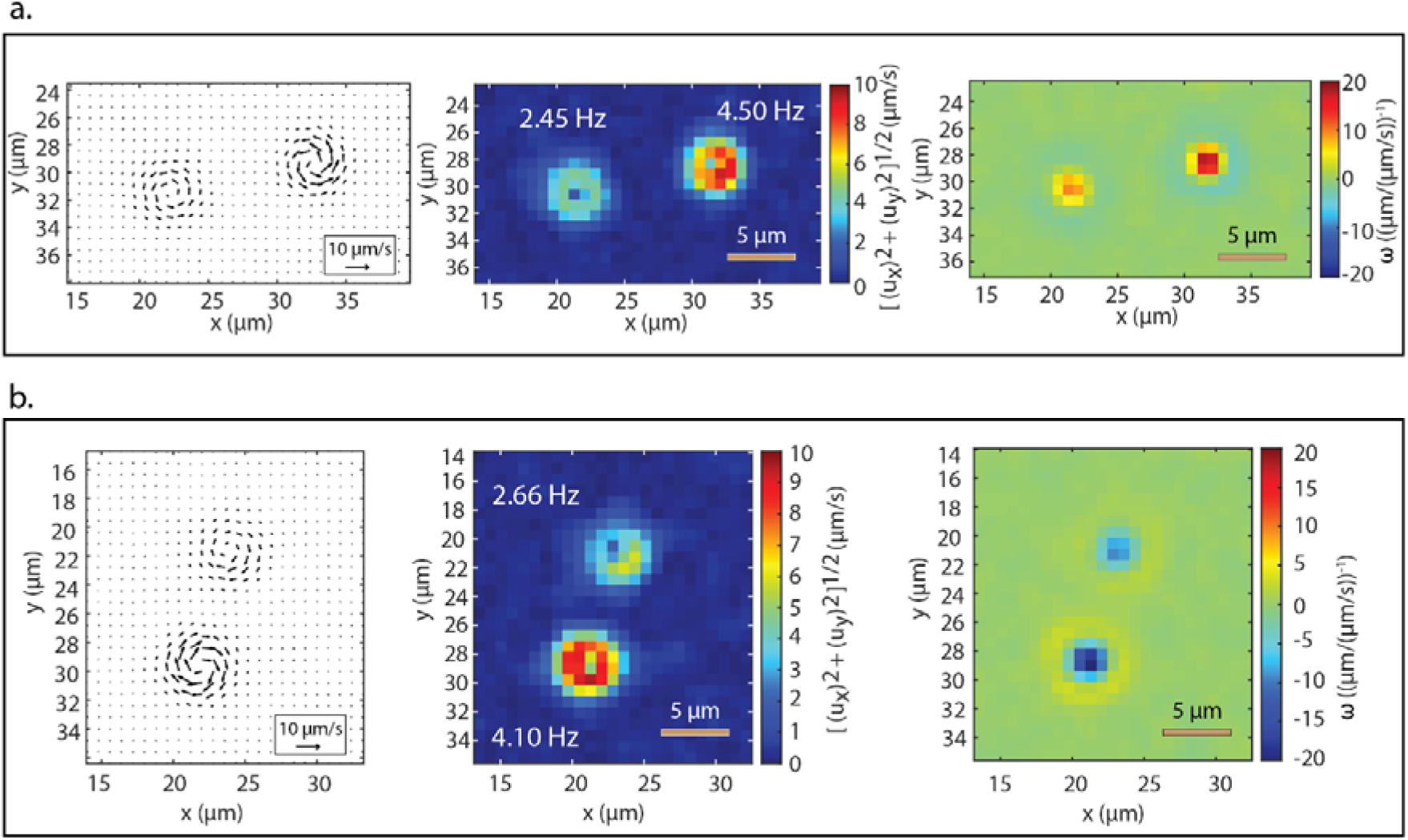
Micro-PIV results of flow fields generated by multiple cell rotations. (a) CCW rotating cells, one is rotating at the speed of 2.45 ± 2.23 Hz and the other rotating at the speed of 4.50 ± 1.48 Hz across x-y plane (25 by 14). (b) CW rotating cells one is rotating at the speed of 2.66 ± 1.16 Hz and the other rotating at the speed of 4.10 ± 0.64 Hz across the x-y plane (20 by 22 μm). Time-averaged velocity arrow (left), velocity magnitude (middle), and vorticity(right).

### Light activated cell rotation

We used CCW rotating bacteria cell where we expressed proteorhodopsin (PR) from a plasmid to control the proton-motive force in response to green light. Initial cell rotation was immediately stopped by 60 mM sodium azide in motility buffer [10 mM KPi (pH 7.0), 85 mM NaCl, 0.1 mM EDTA-Na]. When the sample was illuminated with green laser light (561 nm) at intensity of 55 W/cm^2^, the cell took 30 seconds to start the rotation. Cell speed gradually increased and reached a constant speed of ~ 3.5 Hz after 1-2 minutes (Figure 8). When the green laser was switched off, the cell stopped rotating within 1 millisecond. When the green light was turned on again, the cell started rotating after 1 min and reached a similar constant speed of ~ 3.5 Hz (Figure 8).

**Figure 8.**
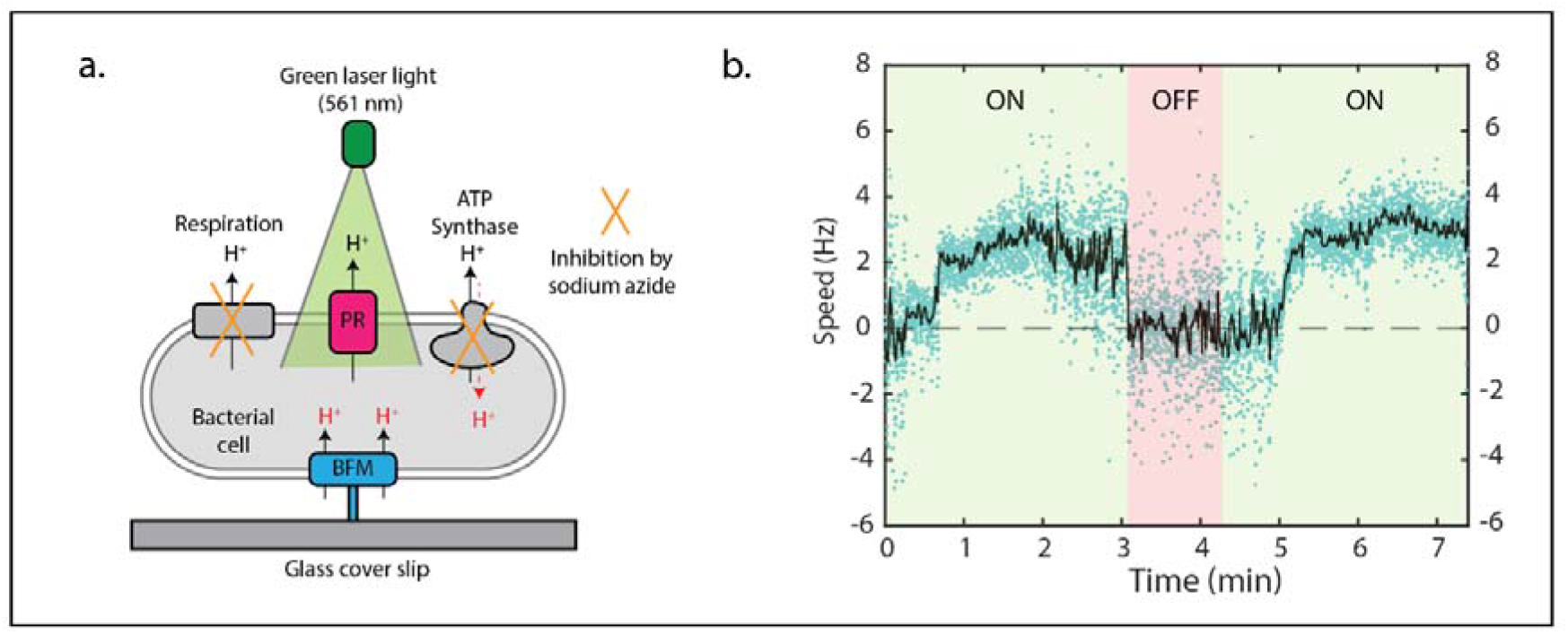
(a) Schematic of PR based green light activated cell rotation via bacterial flagellar motor (BFM). Sodium azide prevents proton pumping outside the cell by inhibiting respiration and ATP synthase function (Walter et al., 2007). (b) Cell rotational speed (black line - 50-point median filtered data, blue dots – original data) versus time for PR expressing CCW rotating cell that was activated by green laser light (561 nm) with the intensity of 55 W/cm^2^. Greenlight switched on (faint green region), and green light switched off (faint red region).

## Discussion

Mixing in stagnant fluids is attained by generating a circular motion of fluid (i.e., vortices) using magnetic stir bars and microspheres. We used the biological rotary motion of micron-sized bacteria to induce mixing in stagnant fluids without the requirement to apply an external electrical or magnetic field. To verify the fluid flows induced by rotation, we used micro-PIV to determine the flow in the fluid caused by bacterial rotation.

Our micro-PIV results showed that single cell bacterial rotation created distinct and localized micro-vortices like magnetic beads that could potentially mix fluids (Ali et al., 2016). We investigated how far into the solution away from the glass coverslip surface the flow could penetrate. We measured that the strength of the flow field decreased exponentially with increasing height from the surface where bacteria were tethered (z-plane). For all the cells we analyzed, flow fields could not be detected at heights above 4 μm. However, this penetration height depended on the cell’s rotational speed. The flow was detected at the z-height of 4 μm for the cell rotating at the speed of 5.75 Hz; by comparison, for the cell rotating at the speed of 3.54 Hz there was no flow detectable at 4 μm above the surface. The geometry of cell tethering could also potentially affect the penetration depth because it is known that rod-shaped bacteria tethered at the center rotate faster than bacteria tethered at the edge (Soman et al., 2020). Likewise, the length of the filament via which the bacterial cell tethers to the surface could affect the penetration depth, as a longer filament would result in a deeper penetration depth. In agreement with the decay with height, the strength of the flow field along the plane of cell rotation (x-y plane) also decreased exponentially with increasing distance from the axis of rotation. The strength of the flow field along the plane was similarly influenced by cell rotational speed, cell length and the size of trace diameter formed by cell rotation (Soman et al., 2020).

We cannot exclude that surface effects from the glass coverslip, from physical contact between the 0.2 μm tracer particle and the 2 μm long cell, or from the shadow of the cell obscuring tracer particle fluorescence might affect the accuracy of our velocity field calculation. We note, however, that the azimuthal velocity 4.6 μm from the centre is 9% of the value 0.8 μm from the centre, that is, the induced vortex persists at least 3 μm beyond the path traced by the arc of the cell (1.66 μm radius). This gives evidence that the velocity effect is observed beyond the boundary of the cell and its direct path. It is not easy to interrogate the space between the cell and the surface, nor is it straightforward to measure the distance between cell and surface for each cell, which can vary. The primary limitation is that it is difficult to integrate tracer particles into this small volume due to particle size and steric hindrance from the cell. Future methods could attempt to tackle this through the use of smaller particles or different fluorescence imaging modalities (noting that high frame rates of ~250 fps are required).

Similarly, it is challenging to simulate boundary effects from the bottom surface since the volume between the cell and the surface has two fixed boundaries in proximity (the cell edge and the surface roughly are 0.5 μm apart). We were able to execute simulations with cells in the centre of a chamber 10 μm in height (5 μm equidistant from the top and the bottom surfaces - SI Figure 8) and observed that at a distance of 1.3 μm above or below the cell the fluid flow velocity had decayed to 21% of the velocity at the plane of the cell. This rules out the possibility of influential boundary effects from the top of our experimental chamber since the top sits ~100 μm above cell. The boundary effect from the coverslip surface is difficult to resolve in our experiments, but what we can observe is that at a height of 4 μm above the cell the flow velocity had decayed to 10% of the velocity in the plane of the cell, and, as above, to 9% at 4.6 μm radially distant from the centre of rotation. Thus we are able to quantify fluid flow around the cell in our tethered cell experiments including potential effects from the near surface.

We note that the average velocity from numerical simulation did not match exactly to our experiment, however both generated circular flow patterns in the surrounding medium with similar maximal velocities and matching trends to our experimental data with the exponential decay of velocity magnitude with increasing z-heights and distance from cell edge in the x-y plane. Our simulations represent an ideal microbial stirbar which rotates at a constant velocity perfectly parallel to the surface. In our experiments, the cell is known to rotate at a varying rotational speed which is normally distributed (Fig. 1d). This fluctuation in rotational speed can be caused by fluctuation in the protonmotive force driving the cell (Gabel & Berg, 2003), or from varying torsional and steric forces that change depending on angle, or a cell that is rotating out of plane, or a tether that is non-centered. We are limited to a cross-sectional slice of the cell when measuring the dimensions of the cell, and we do not know the exact tether location or path through which the cell moves. Nevertheless, we initialized our simulations using matched geometries for the cells and measurements of the cell arc, which closely matched tethered cell geometries and speeds from our experiments (Islam et al., 2020, 2022) and others’ (Soman et al., 2020), and saw comparable decay in velocity magnitude with distance from the cell in all directions.

When mixing solutions using a rotating object, control over the direction of rotation and the surface patterning of the rotating object are important for the development of efficient micromixers (J.-W. Kim & Tung, 2015). In our work, we have demonstrated directional control over cell rotation through the genomic engineering of bacteria to rotate in CCW or CW direction. We showed that rotation of single cell generated flow with localized micro-vortices and that these vortices’ directionality corresponded to the rotation direction. The rotation of multiple cells also caused distinct local micro-vortices for each cell that was rotating. We saw no collective behavior due to hydrodynamic interaction in the region between two vortices generated by unsynchronized cells separated by ~10 μm. We cannot rule out vortex interaction at closer spacing, as we did not record data where the cells were more densely packed. While we can easily generate very sparse cell densities as we can control the concentration of the cells and the strength of the surface interaction, it is challenging to generate periodic arrays of regularly spaced rotating cells as the adhesion of the tether to the surface is still stochastic and at higher surface densities of cells the cells can sterically hinder the rotation of other cells. Work with smaller cells, or smaller particles, or alternate imaging modalities may allow testing of vortex interaction at closer spacings. Our findings agree with previous work which held the separation distance between two rotating microspheres at ten times the diameter of the microspheres and observed independent vortices with unresolvable flow patterns in the region between the microspheres (Ali et al., 2016).

Coordination and correlation of tethered micromixers may require cell patterning at a periodic and near spacing to amplify the hydrodynamic interaction. Furthermore, synchronization of rotation may be useful. In our experiments, we relied on the random settlement of bacteria to tether on the surface and did not have control over the tethering of multiple bacterial cells to induce tethering at repeat and near distances. Other properties that varied among the bacterial population were the rotational speed, cell length, and geometry of cell tethering (Supplementary Video 1). To pattern cells with high spatial control many techniques may offer future avenues: for example, optical trapping, cell confinement wells and microfabricated sieves have been used to tether cells with high localization on the substrate (J.-W. Kim & Tung, 2015).

External control of mixing is desirable when attempting to mix solutions in confined structures in microfluidic systems. We demonstrated that mixing could be controlled remotely by switching on and off the rotation of bacterial cells using light. We used green light to activate the rotation of bacteria by expression of the light sensitive proton pump PR which can pump protons to generate proton motive force when illuminated. Magnetic fields offer another method of external control; however, use of light offers several advantages in terms of cost, portability, and its ease of use in complex but transparent microfluidics designs (Venancio-Marques et al., 2013).

We observed that light activation via restoration of proton motive force from PR expression took at least 30 seconds to activate rotation upon illumination with green light. Future work could decrease this activation time to trigger mixing more rapidly. In this work, we applied sodium azide to stop the cell rotation, and this is toxic to human cells (Tat et al., 2021). Use of sodium azide could be substituted by deleting the gene encoding ATP synthase to switch off the cell rotation (Arlt et al., 2018). However, it was still necessary to wait up to 10 minutes until the oxygen in the solution was consumed stopping cell rotation (Arlt et al., 2018). Future work could explore more direct methods of light activation; for example, engineering stators to directly incorporate a light activatable components (Zhang et al., 2020), thus, decoupling the destruction and restoration of proton-motive force with the light activation of rotation, which necessarily places a higher burden on the cell.

In conclusion, we have demonstrated that bacterial cell rotation can be employed for mixing stagnant fluids in microfluidics at low Reynolds number. Rotation of bacterial cell bodies generated distinct micro-vortices with circular fluid motion which degenerated with increasing height and distance away from the center of rotation. We further demonstrated that cell rotation could be remotely activated using laser illumination, thus providing external control to switch on and off the rotation for mixing application. Besides the applications in mixing, bacterial motility could also be used to power micropumps (Dauparas et al., 2018), as a power generator (Tung & Kim, 2006), and for biosensors (Krasnopeeva et al., 2020). In future, various challenges such as specific tethering of cells, synchronized cell rotation, longer cell viability, and minimizing varying cell speed are necessary to be addressed to establish bacterial cell rotation as an efficient micro mixer. Furthermore, genetic engineering of bacterial flagellar motor with new light-sensitive domains could be beneficial to activate the mixing.

## Materials and methods

### Sample preparation

In this paper, two types of bacterial strains were used, both with FliC-sticky filaments. One of the bacterial strains rotated in counterclockwise direction (RP437 *ΔmotAB ΔcheY fliC*^st^ (Sowa et al., 2005)with MotAB expressed from plasmid, pDB108) Whereas the other strain rotated in clockwise direction (RP437 *ΔfliG fliC^st^* with FliG-dPAA expressed from plasmid (Minamino et al., 2011)). Both the strains were transformed with plasmid encoding green fluorescent protein (pACGFP1) for visualization, thus, recording speed and length of rotating cell. Cell tethered experiment was performed in glass-cover slip tunnel slide (Ishida et al., 2019). For multiple cell tethering, bacteria (OD_600_ 0.5-0.6) were incubated for 15 min to settle down to the surface of tunnel slide whereas for single cell, bacteria (OD600 0.5-0.6) were incubated for 5 min.

Likewise for the light activation experiment, sticky filament bacterial strain was transformed with a plasmid encoding PR (Arlt et al., 2018). OD_600_ of bacterial sample was prepared as previously mentioned with addition of 10 μM all-trans-Retinal in the culture media. Before activating with light, bacterial sample was suspended in motility buffer supplemented with 60 mM of sodium azide to stop the cell from rotating (Walter et al., 2007) to create a situation where cells are not rotating as an initial condition prior to light activated rotation. A full list of strains and plasmids is shown in Supplementary Table 1.

### Micro-PIV experiment

A tunnel slide containing tethered bacteria and red fluorescent microspheres (200 nm diameter) with seeding density of 4.6 x 10^12^ particles/mL was mounted in the fluorescent microscope (Figure 1a-c). A 642 nm laser was used to illuminate red fluorescent tracing microspheres whereas 488 nm laser was used to illuminate GFP expressing bacterial cell. Total internal reflection fluorescence (TIRF) mode to investigate flow field near the surface whereas highly inclined and laminated optical sheet (HI-Lo) mode to investigate at different z-heights. For each experiment, two sets of videos were recorded using sCMOS camera in the same field of view. One video of rotating green, fluorescent tethered cell with 500 frames was recorded for 12 seconds (~ 42 fps) whereas another video of red fluorescent beads with 5000 frames was recorded for 23 seconds (~ 217 fps) as micro-PIV data acquisition. Rotational speed of cell was analyzed using LabView software and cell length was determined using ImageJ software. Thus acquired PIV data was processed using commercial software package (DaVision 8.4) with three passes of 16 by 16-pixel interrogation window (50% overlap). Average velocity of image sequences was estimated based on time-averaged correlation method by using open-source MATLAB based PIVmat toolbox version 4.20 (Moisy F., 2023) and visualized as graphical plots using Prism GraphPad software. Time averaged velocities averaged across all frames are shown in Figs 2, 5, & 7.

### Light activation of cell rotation

For external activation of cell rotation, a green laser of 561 nm wavelength was used to illuminate both the cells and microspheres. Upon green light absorption by PR, tethered bacterial cells start to rotate in response to the decreased concentration of proton inside the bacterial cell by PR activation. To visualize the cells, white light was passed through long-pass RG630 filter to prevent excitation of PR during visualization.

### Numerical simulation

Finite element computational simulations using the Fluid Flow (FLUENT) module of ANSYS were conducted to capture the velocity fields caused by rotating a microbial stir bar at the X-Y plane at various cross-sections along the Z-axis. SI Figure 6a show the reconstructed bacterial cell for the simulation, and SI Figures 6b and 6c shows the cell placed parallel to a surface inside the confined computational environment with a small gap between the bottom of the cell and the underlying surface, and the top view cross-section of the modelled system, respectively.

The multi-zone unstructured tetrahedral computational mesh was generated with a mesh size chosen based on a mesh independent analysis with orthogonal quality of 0.78 and an average skewness of 0.21. Dynamic meshing with smoothing was used to simulate the motion of the cell. To simulate the rotation of the cell, a sliding wall boundary condition with a predefined angular velocity vector was applied to the cell’s surface. An adiabatic no-slip boundary condition, where the velocity is zero, was used at the walls in computational geometry.

Simulations were conducted to solve the incompressible form of the three-dimensional transient Navier–Stokes equations, including momentum and continuity equations. The fluid’s properties surrounding the cell were chosen to match those of water at room temperature. The fluid was considered an isothermal static liquid. The SST k-omega turbulence model was used to capture the turbulent flow regime. The Semi-Implicit Method for Pressure Linked Equations (SIMPLE) algorithm was used for pressure velocity coupling, and the standard algorithm was employed for pressure interpolation. The first-order upwind differencing was used for the spatial discretization of transport equations. In each time step (1 ms), a maximum of 10 iterations was found to be sufficient for convergence, and all solutions converged with residuals below 10E-6. Simulation figures show transient velocities at final time step of simulation (Fig. 6).

## Supplementary Material

See Supplementary Material for Supplementary Figures 1-11, Supplementary Table (of primers) and Supplementary Videos showing cell rotation.

## Acknowledgements

This work was conducted (in part) using the facilities at the South Australia node of the Australian National Fabrication Facility (ANFF-SA), a company established under the National Collaborative Research Infrastructure Strategy to provide nano- and microfabrication facilities for Australia’s researchers. MABB acknowledges funding support from a Scientia Fellowship from UNSW, Human Frontiers Science Program Project Grant RGY0072/21 and US Navy Office of Naval Research Research Grant N62909-22-1-2051.

## Conflict of interest

There are no conflicts to declare.

## Author contributions

JPG and MABB executed and designed experiments in molecular biology, microfluidics and light activation. MNK, JPG and MABB designed simulation framework to match experimental measurements. MNK executed numerical simulations. JPG, CMdS and MABB designed and executed experiments and analysis for particle image velocimetry. CMdS supervised the analysis of particle image velocimetry experiments. MABB supervised the design, execution and writing of the project. All authors contributed to the writing and revision of the manuscript.

## Notes

### Competing Interest Statement

The authors have declared no competing interest.

### Summary of Updates

We have added additional Supplementary Figures for simulation data and more discussion on limitations.

