## Supplementary Figures 1-11, Supplementary Table 1 for "Microbial stir bars: light-activated rotation of tethered bacterial cells to enhance mixing in stagnant fluids"

**Supplementary Material**

**This PDF file includes:**

Supplementary Note

Supplementary Figures 1 to 11

Supplementary Table 1

Captions for Supplementary Videos 1 to 9

**Other Supplementary Materials for this manuscript includes the following:**

Supplementary Videos 1 to 9

**Supplementary Note**

**Equation for the depth of field (δz) calculation.**

The following equation was used to calculate the depth of field for μ-PIV experiment (Kim et al., 2015):

$$\delta z = \frac{3n\lambda}{NA^2} + \frac{2.16d_p}{\tan\theta} + d_p$$

where 'n' is the refraction index of immersion oil used between coverslip and objective lens, 'λ' is laser wavelength, 'NA' is numerical aperture of our objective lens, '$d_p$' is the diameter of fluorescent tracing particles, and 'Θ' is small light collection angle. In our case, n = 1.518, λ = 0.633 μm (633 nm), NA = 1.46, $d_p$ = 0.2 μm (200 nm), Θ = 74.11°. Thus, depth of field (δz) was calculated as 1.67 μm.

**Supplementary Figures**

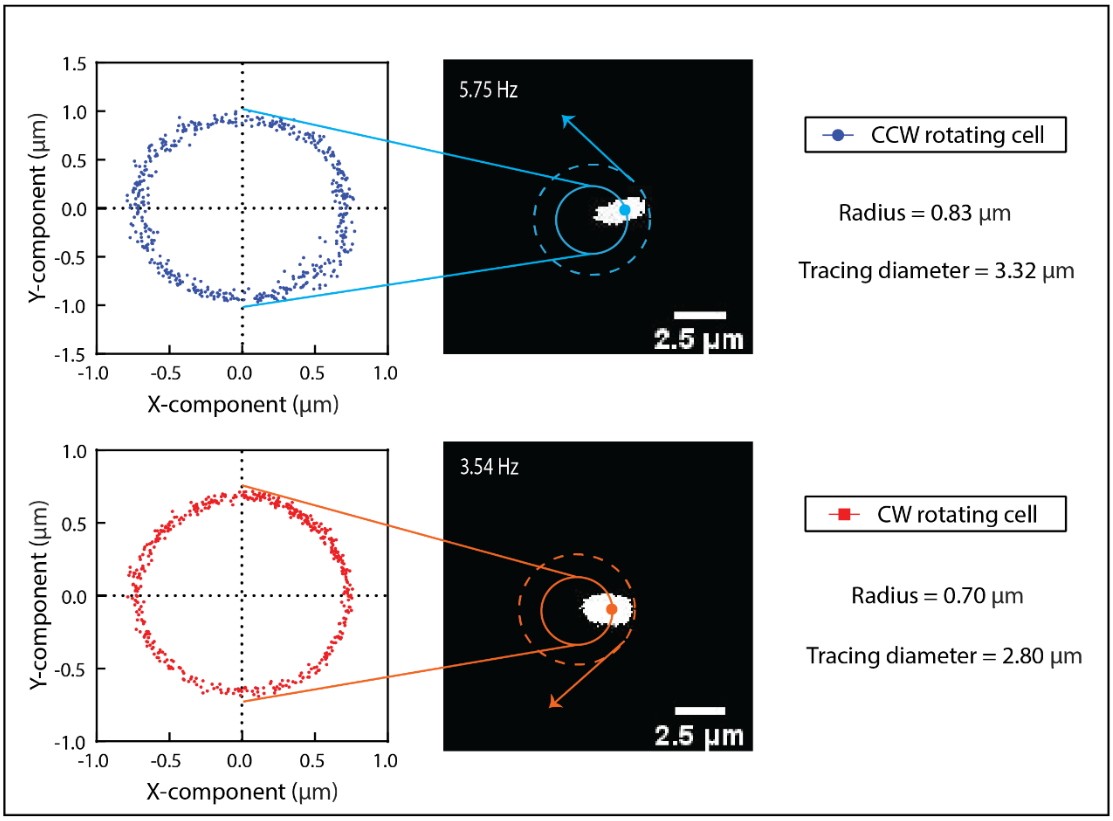

**SI Figure 1. Rotational traces of cell centroids.** Trace diameter measurement of CCW rotating cell (5.75 ± 0.39 Hz), represented by blue scatter plot and CW rotating cell (3.54 ± 0.58 Hz), represented by red scatter plot. Individual data points for X and Y component of cell centroid traversed by rotating cell were fitted by circle. The radius of fitted circle was multiplied by 4 to obtain the tracing diameter.

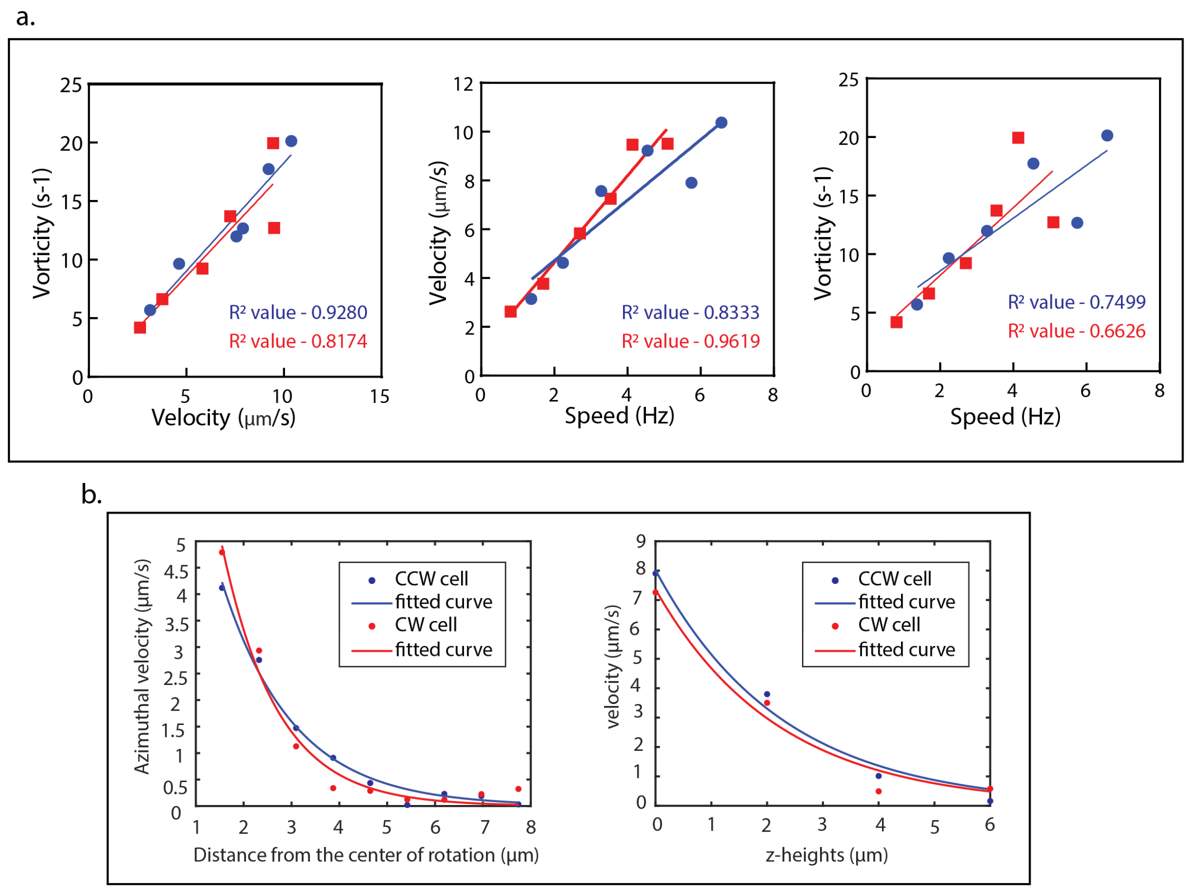

**SI Figure 2. Correlation of flow speed with rotational cell speed and decay with distance from rotational mixer.** Curve fitting using simple linear regression model by equation y = m*x + c, where 'y' is velocity or vorticity, 'x' is speed, 'm' is the slope and 'c' is the y-intercept. First-term exponential fit using equation $y = a*e^{b*x}$, where 'y' is velocity, 'a' is initial velocity, 'b' is velocity decay rate/constant, and 'x' is z-heights or distance from the center of the rotation. (a) Linear fit for vorticity vs velocity, velocity vs speed, and vorticity vs speed with their respective $R^2$ value. Red-color filled square, and line represent CW rotating cell and blue-color filled circle and line represent CCW rotating cell. (b) First-term exponential fit for azimuthal velocity vs distance from the center of rotation and maximum velocity vs z-heights. Blue-color filled circle represents original data points whereas blue line represents exponentially fitted curve for CCW rotating cell (5.75 ± 0.39 Hz). Fitted curve, represented by the equation, $y = 11.93*e^{-0.67*x}$ (velocity vs distance from the center of rotation) and $y = 8.02*e^{-0.4431*x}$ (velocity vs z-heights) with $R^2$ values of 0.99 and 0.98 respectively. The red color filled circle represents original data points whereas red line represents exponentially fitted curve for CW rotating cell (3.54 ± 0.58 Hz). Fitted curve, represented by the equation, $y = 18.72*e^{-1.06*x}$ (velocity vs distance from the center of rotation) and $y = 7.36*e^{-0.45*x}$ (velocity vs z-heights) with $R^2$ values of 0.98 and 0.97 respectively.

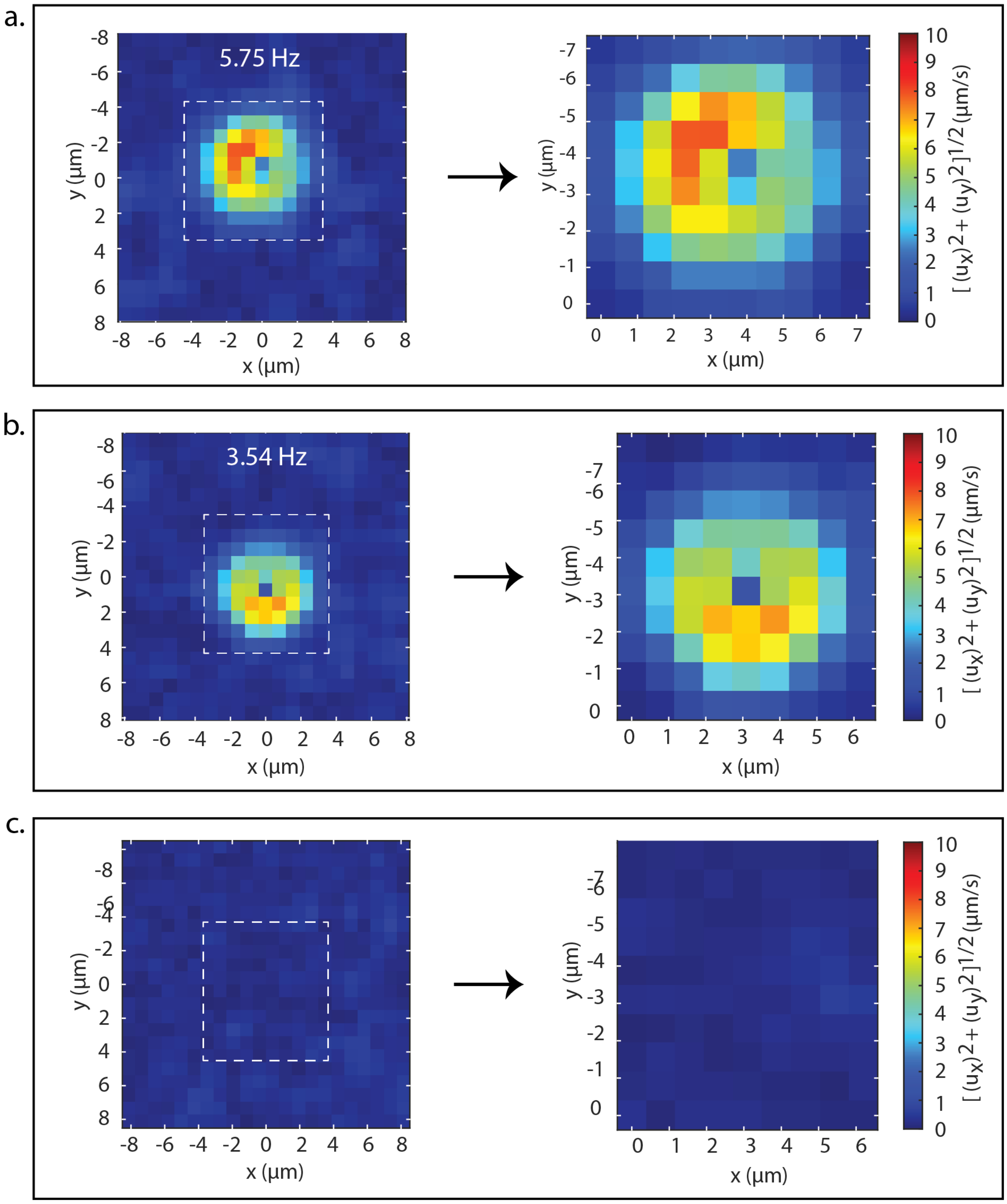

**SI Figure 3. Comparison of rotating and non-rotating regions of interest.** Region of interest extracted for time-averaged velocity magnitude calculation, shown in right panel. (a) CCW rotating cell at the speed of 5.75 ± 0.39 Hz across x-y plane (8 by 8 μm). (b) CW rotating cell at the speed of 3.54 ± 0.58 Hz across x-y plane (7 by 8 μm). (c) Sample without any rotating cells i.e., with only fluorescent particles as a negative control across x-y plane (7 by 8 μm).

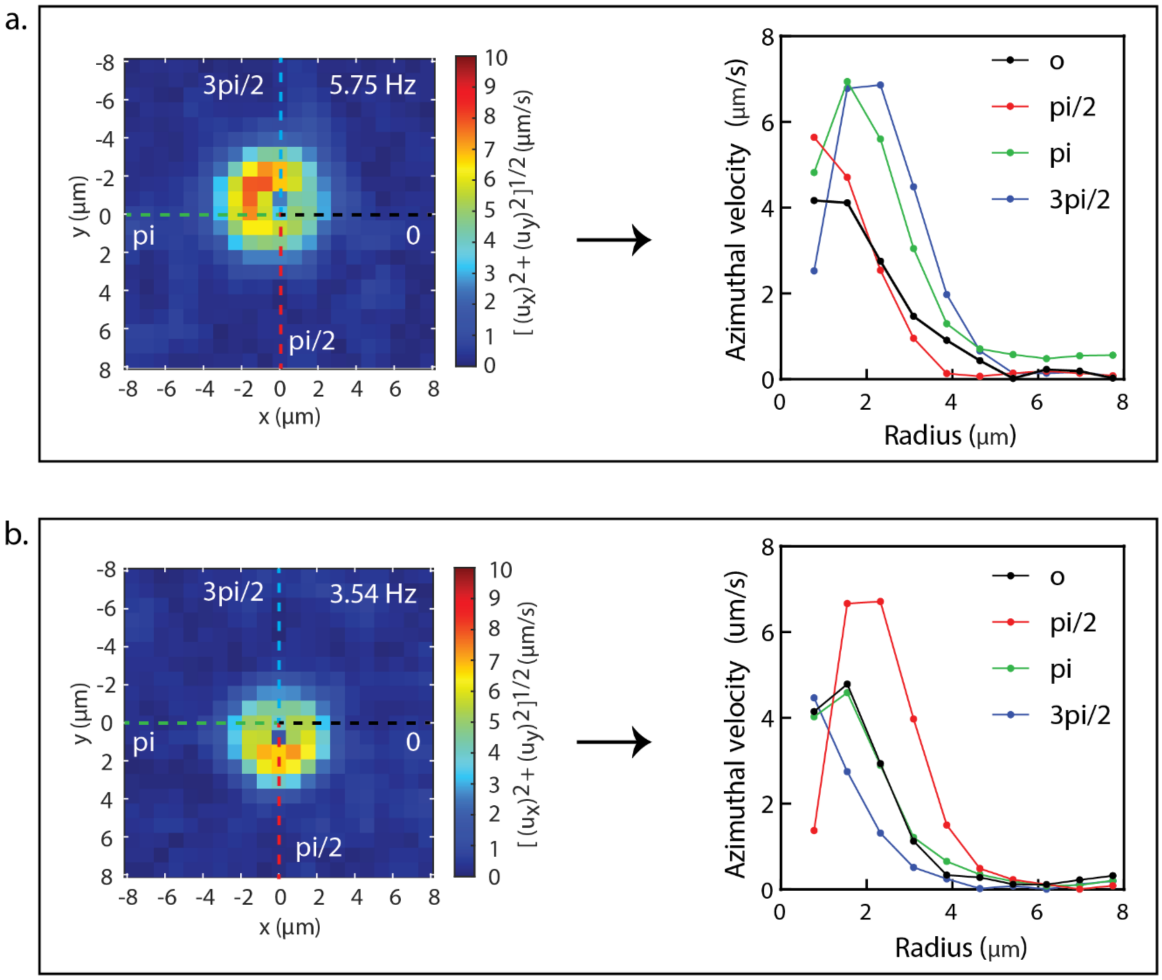

**SI Figure 4. Decay in tangential velocity with distance from the centre of the rotational mixer.** Azimuthal (tangential) velocity along the radius passing through array of angles sampled between 0 and 2*pi in x-y plane. Left panel – velocity magnitude plot shown with dotted line with different colors for the radius passing through respective angles. Right panel – plot of azimuthal velocity vs radius or distance away from the center of rotation. (a) CCW cell rotating with speed of 5.75 ± 0.39 Hz. (b) CW cell rotating with the speed of 3.54 ± 0.58 Hz.

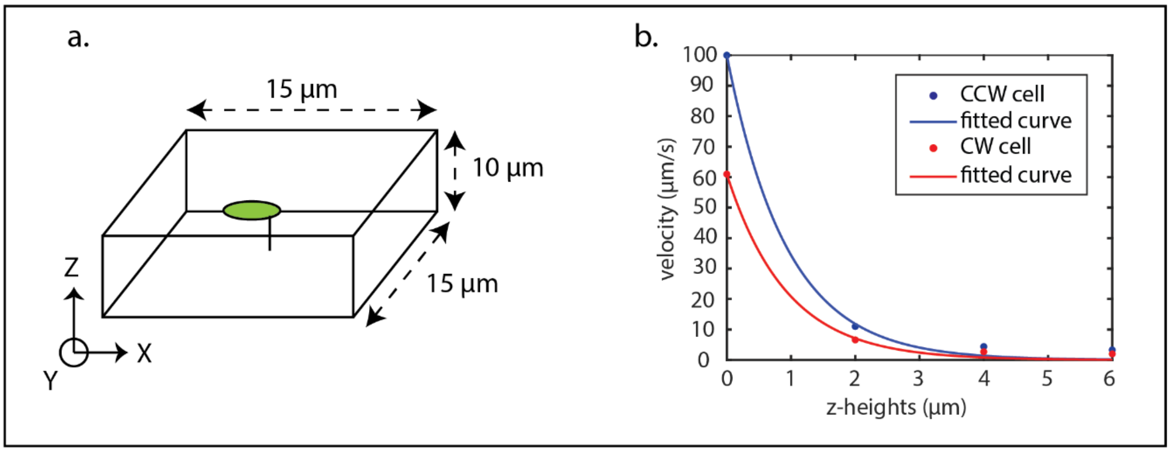

**SI Figure 5. Simulated decay in velocity with distance from cell.** (a) Schematic diagram of geometrical structure for CFD simulation, green elongated cell rotating in CCW or CW direction inside the rectangular box with length of 15 µm, width of 15 µm and height of 10 µm, with cell located in the centre of the chamber at a height of 5 µm. (b) First-term exponential fit of maximum velocity vs z-heights for simulated data. First-term exponential fit using equation $y = a*e^{b*x}$, where' y' is velocity, 'a' is initial velocity, 'b' is velocity decay rate/constant, and 'x' is z-heights or distance from the center of the rotation. Blue-color filled circle represents original simulated data whereas blue line represents exponentially fitted curve ($y = 99.95*e^{-1.066*x}$) for CCW rotating cell (5.75 Hz) with $R^2$ value of 0.99. The red color filled circle represents original simulated data whereas red line represents exponentially fitted curve ($y = 60.97*e^{-1.074*x}$) for CW rotating cell (3.54 Hz) with $R^2$ value of 0.99.

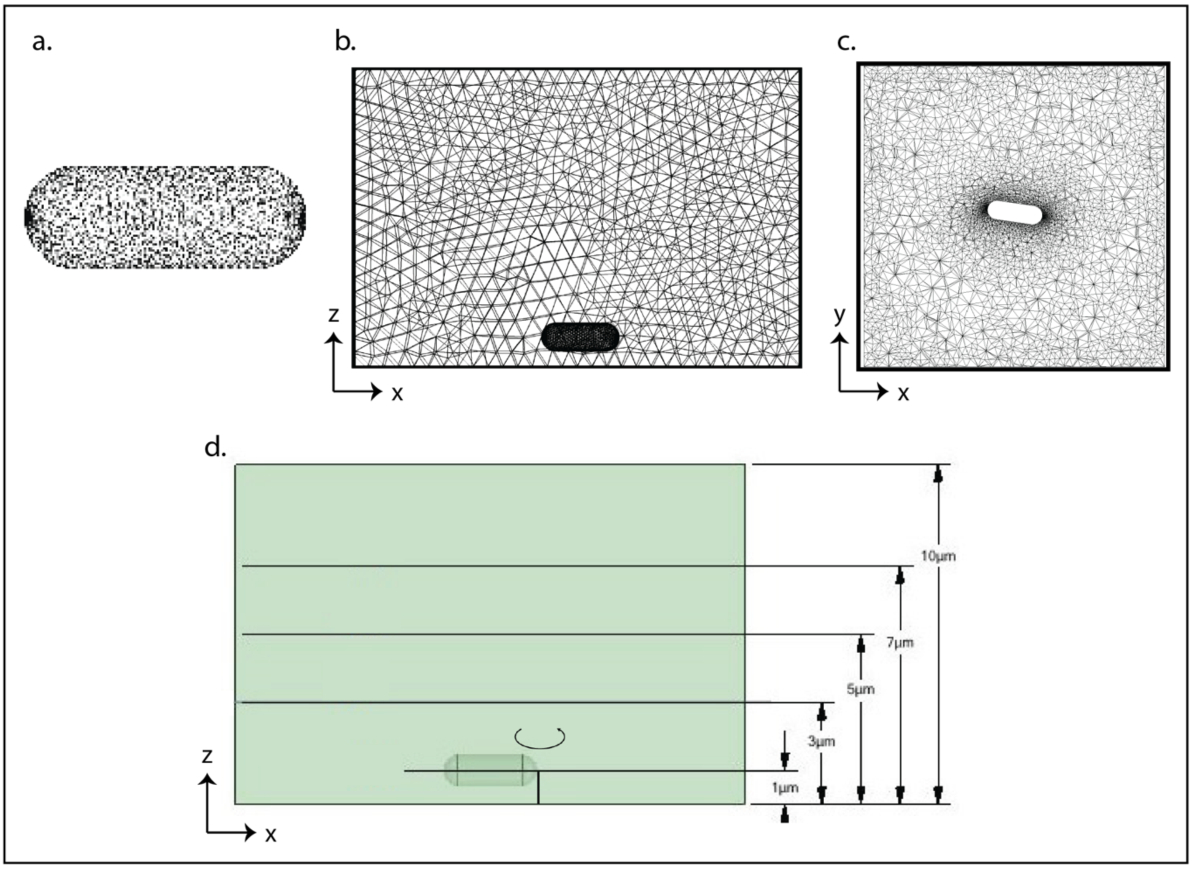

**SI Figure 6. Mesh used for simulations.** (a) the reconstructed discretized in the computational elements, (b) the cell parallel to a surface of the computational domain with a small opening between the bottom of the cell and the underlying surface. Box is 10 μm in z and 15 μm in x. (c) Top view cross-section of the system. Box is 15 μm in x by 15 μm in y. (d) Geometry of the system, with cell rotating with central plane 1.0 μm from the surface (edge of cell 0.5 μm from surface) and simulation cut-slices calculated at 2 μm, 4 μm and 6 μm above the central plane of the cell.

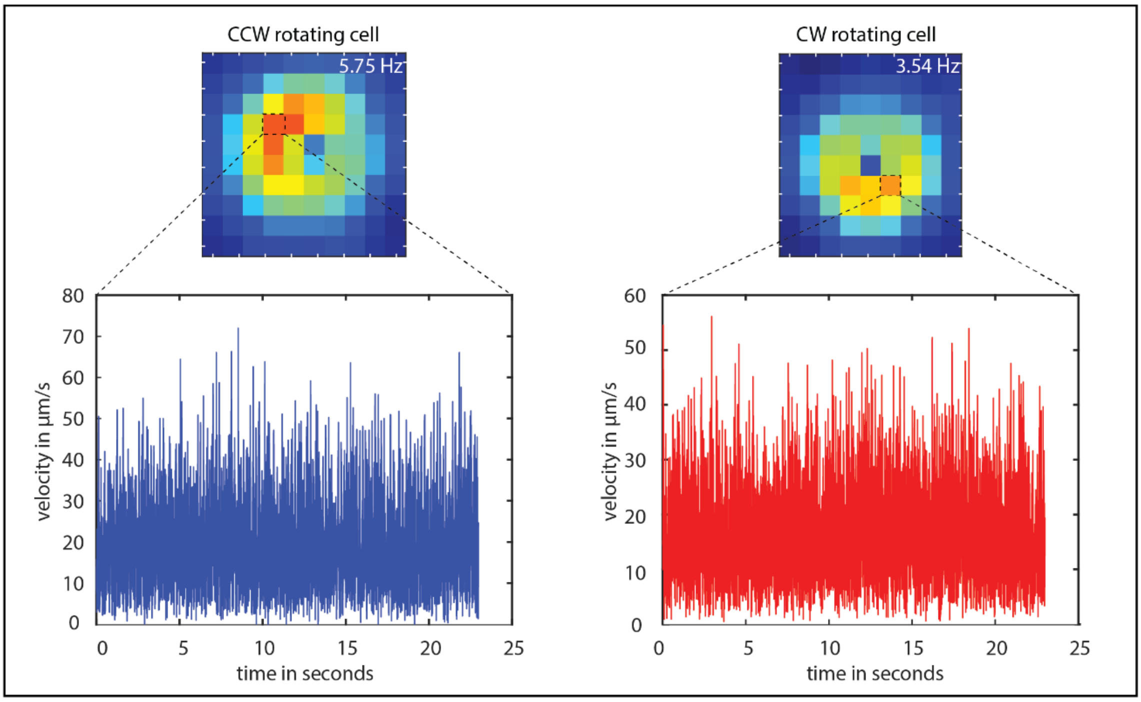

**SI Figure 7. Time evolution and fluctuation of maximum velocity for a given spatial domain.** Maximum velocity is calculated across a 16 x 16 pixels of interrogation window, where maximum velocity pixel is indicated by dotted square box. (Left) CCW rotating cells at the speed of 5.75 ± 0.39 Hz with maximum velocity of 71.96 μm/s. (Right) CW rotating cells at the speed of 3.54 ± 0.58 Hz with maximum velocity of 56.11 μm/s.

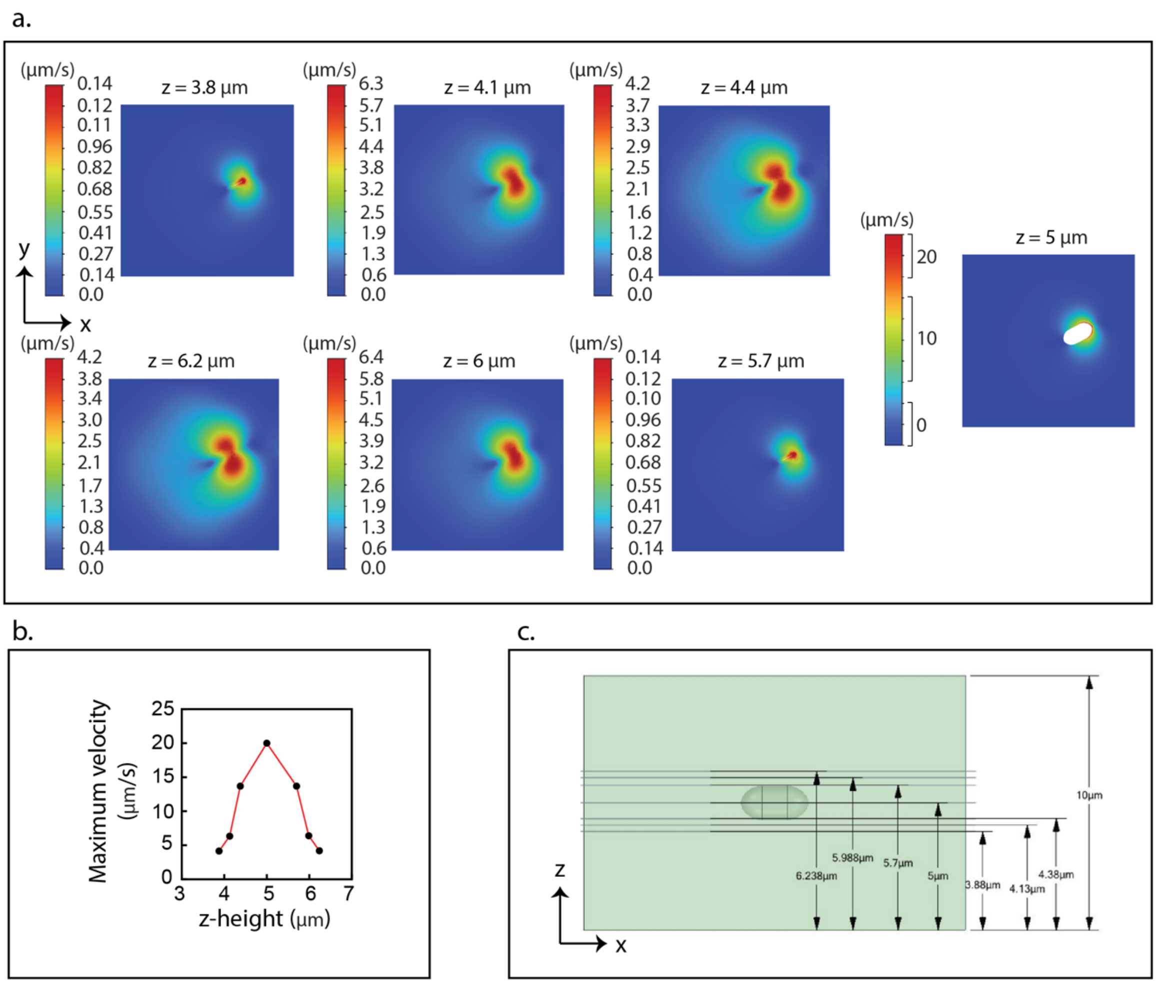

**SI Figure 8. Numerical simulation of instantaneous velocity magnitude generated by single-cell rotation at different z-heights.** (a) Simulation of velocity generated in the spatial domain above and below the cell rotation plane i.e., distance from bottom surface**.** The cell used for simulation was a CCW rotating cell with the speed of 1.13 Hz rotating in the x-y plane (15 by 15 µm) for 1.13 seconds. (b) Plot of maximum velocity vs z-height showing overall fluid velocity response in z away from the cell. (c) Geometry of the simulation. Cell was positioned in the centre of the chamber at a z-height of 5 µm. Each cell was identical in shape, defined by the length of 2.75 µm and width of 1.5 µm. Cell was positioned in the centre of the chamber at a z-height of 5 µm.

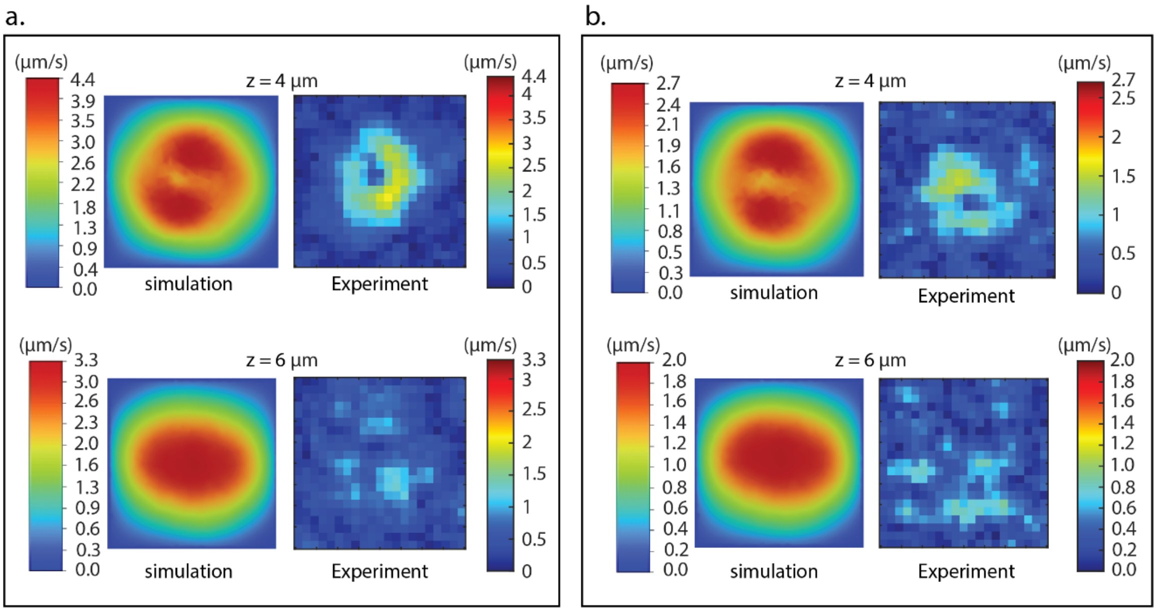

**SI Figure 9. Comparison between instantaneous velocity magnitude from numerical simulation and time-averaged velocity magnitude from micro-PIV experiment for single-cell rotation at 4 and 6 μm in z-height.** (a) CCW rotating cell with the speed of 5.75 Hz across the x-y plane (15 by 15 μm) at 0.175 seconds (simulation) and time-averaged over 23 seconds (experiment). (b) CW rotating cell with the speed of 3.54 Hz across the x-y plane (15 by 15 μm) at 0.301 seconds (simulation) and time-averaged over 23 seconds (experiment). Both cells were elongated in rod like shape with axis of rotation at the edge, defined by a length of 2.75 μm and a width of 1.0 μm. Simulations were run for one full revolution of the cell with a timestep of 1 ms.

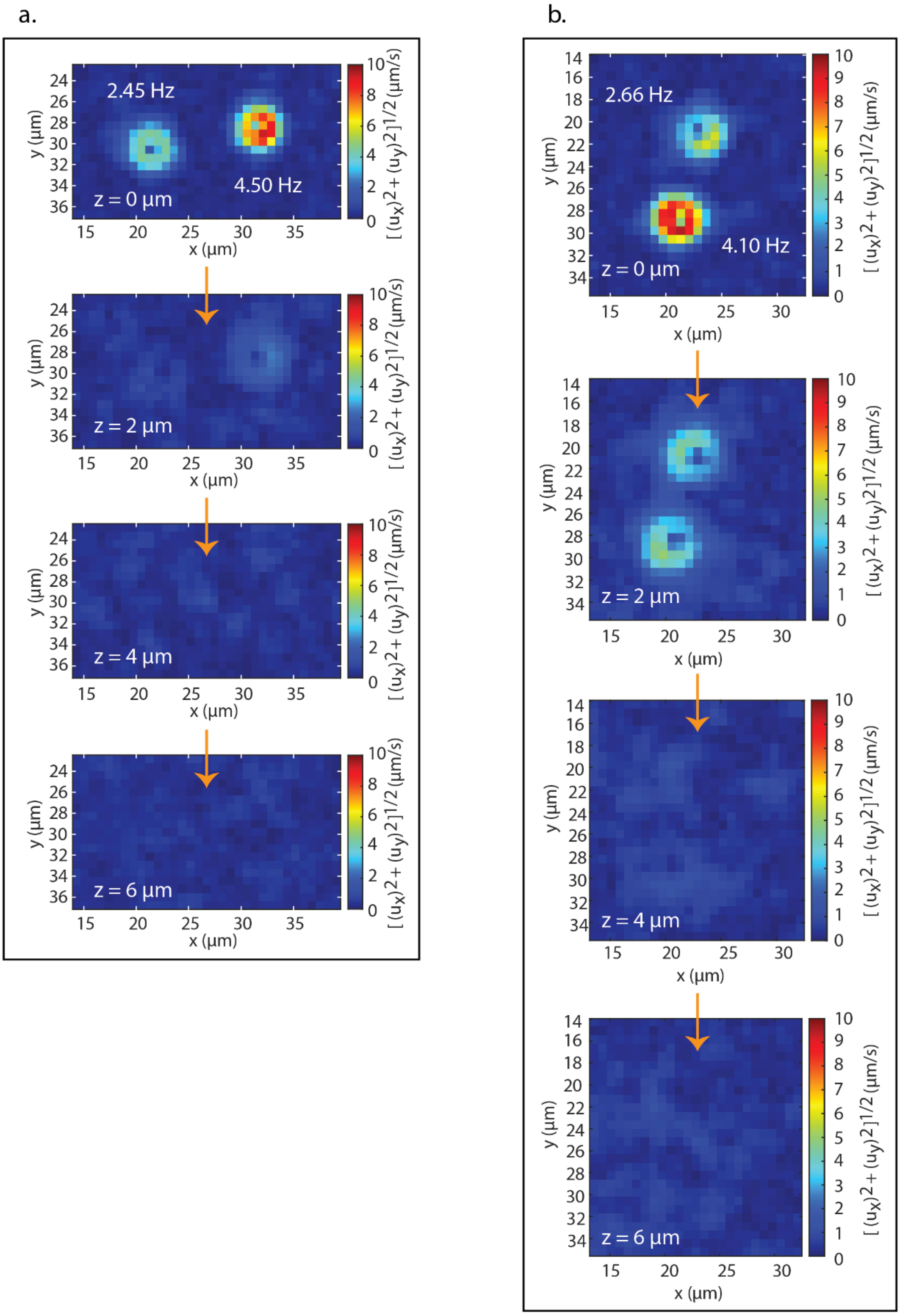

**SI Figure 10. Time-averaged velocity magnitude measured for multiple cell rotations at different z-heights.** (a) CCW rotating cells, one rotating at the speed of 2.45 ± 2.23 Hz and other rotating at the speed of 4.50 ± 1.48 Hz across x-y plane (25 by 14) at 0, 2, 4, and 6 μm of z-heights (top to bottom). (b) CW rotating cells one rotating at the speed of 2.66 ± 1.16 Hz and other rotating at the speed of 4.10 ± 0.64 Hz across x-y plane (20 by 22 μm) at 0, 2, 4, and 6 μm of z-heights (top to bottom).

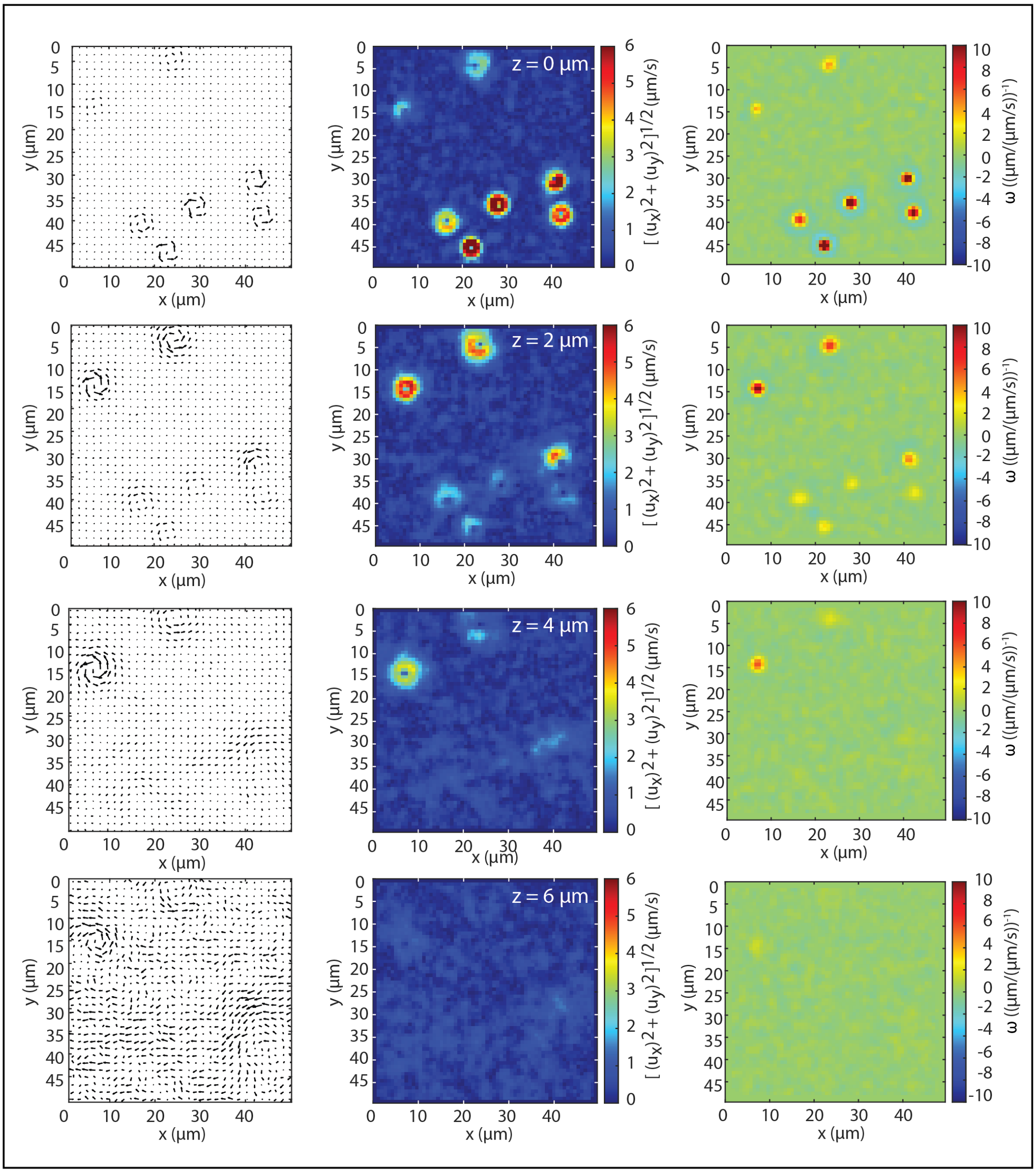

**SI Figure 11. Time-averaged velocity arrow, velocity magnitude, and vorticity plot for multiple CCW rotating cells (more than two cells).** Fluid flow analyzed at different z-heights of 0, 2, 4, and 6 µm (top to bottom).

**Supplementary Table 1**. **List of strains and plasmids used for the experiment.**

ARA, Arabinose; CAM$^R$, Chloramphenicol resistant, AMP$^R$, Ampicillin resistant

| **Strains** | **Description** | **Reference** |
|---|---|---|
| ΔFliG-S | *E. coli* RP437 (ΔFliG, FliC-sticky) | (Minamino et al., 2011) |
| SYC-38 | *E. coli* RP437 (ΔMotA, ΔMotB, ΔCheY, FliC-sticky) | Yoshiyuki Sowa |
| **Plasmids** | **Description** | **Reference** |
| pDB108 | MotA and MotB, CAM$^R$ | David F Blair |
| p-dPAA-FliG | FliG, CAM$^R$ | (Minamino et al., 2011) |
| pBAD-HisC-PR | Proteorhodopsin (PR), AMP$^R$ | (Arlt et al., 2018) |
| pACGFP1 | Green Fluorescent Protein (GFP), AMP$^R$ | (Dong et al., 2016) |

**Captions for Supplementary Videos.**

**Supplementary Video 1.** Bright field microscopic view of multiple cells rotating in CCW direction, surrounded by buffer containing dark red fluorescent microspheres (0.2 μm in diameter). Videos are presented with 30 fps frame rate.

**Supplementary Video 2.** CFD simulated contours of velocity magnitude from time 0.011 s to 0.175 s for CCW rotating cell (5.75 Hz), situated at the z-height of 0 μm.

**Supplementary Video 3.** CFD simulated contours of velocity magnitude from time 0.011 s to 0.175 s for CCW rotating cell (5.75 Hz), situated at the z-height of 2 μm.

**Supplementary Video 4.** CFD simulated contours of velocity magnitude from time 0.011 s to 0.175 s for CCW rotating cell (5.75 Hz), situated at the z-height of 4 μm.

**Supplementary Video 5.** CFD simulated contours of velocity magnitude from time 0.011 s to 0.175 s for CCW rotating cell (5.75 Hz), situated at the z-height of 6 μm.

**Supplementary Video 6.** CFD simulated contours of velocity magnitude from time 0.011 s to 0.301 s for CW rotating cell (3.54 Hz), situated at the z-height of 0 μm.

**Supplementary Video 7.** CFD simulated contours of velocity magnitude from time 0.011 s to 0.301 s for CW rotating cell (3.54 Hz), situated at the z-height of 04μm.

**Supplementary Video 8.** CFD simulated contours of velocity magnitude from time 0.011 s to 0.301 s for CW rotating cell (3.54 Hz), situated at the z-height of 4 μm.

**Supplementary Video 9.** CFD simulated contours of velocity magnitude from time 0.011 s to 0.301 s for CW rotating cell (3.54 Hz), situated at the z-height of 6 μm.
